# Accurate prediction of cohesin-mediated 3D genome organization from 2D chromatin features

**DOI:** 10.1101/2022.11.23.517572

**Authors:** Ahmed Abbas, Khyati Chandratre, Yunpeng Gao, Jiapei Yuan, Michael Q. Zhang, Ram S. Mani

## Abstract

The three-dimensional (3D) genome organization influences diverse nuclear processes. Chromatin interaction analysis by paired-end tag sequencing (ChIA-PET) and Hi-C are powerful methods to study the 3D genome organization. However, ChIA-PET and Hi-C experiments are expensive, time-consuming, require tens to hundreds of millions of cells, and are challenging to optimize and analyze. Predicting ChIA-PET/Hi-C data using cheaper ChIP-Seq data and other easily obtainable features could be a useful alternative. It is well-established that the cohesin protein complex is a key determinant of 3D genome organization. Here we present Chromatin Interaction Predictor (ChIPr), a suite of regression models based on deep neural networks (DNN), random forest, and gradient boosting, respectively, to predict cohesin-mediated chromatin interaction strength between any two loci in the genome. Comprehensive tests on four cell lines show that the predictions of ChIPr correlate well with the original ChIA-PET data at the peak-level resolution and bin sizes of 25 and 5 Kbp. In addition, ChIPr can accurately capture most of the cell-type-dependent loops identified by ChIA-PET and Hi-C data. Rigorous feature testing indicated that genomic distance and RAD21 (a cohesin component) ChIP-Seq signals are the most important inputs for ChIPr in determining chromatin interaction strength. The standard ChIPr model requires three experimental inputs: ChIP-Seq signals for RAD21, H3K27ac (enhancer/active chromatin mark) and H3K27me3 (inactive chromatin mark). The minimal ChIPr model performs comparably and requires a single experimental input: ChIP-Seq signals for RAD21. Integrative analysis revealed novel insights into the role of CTCF motif, its orientation, and CTCF binding on the prevalence and strength of cohesin-mediated chromatin interactions. These studies outline the general features of genome folding and open new avenues to analyze spatial genome organization in specimens with limited cell numbers.

## INTRODUCTION

The three-dimensional (3D) genome organization directly impacts diverse nuclear processes such as transcription, DNA repair, and replication. Therefore, it is crucial to understand how the distal regulatory elements (in the linear genome) interact in 3D space. Several sequencing-based and imaging-based experimental methods have been developed in the last two decades to study the 3D genome organization ^1^. Many of the sequencing-based approaches are derived from the chromosome conformation capture (3C) concept ^2^. High-throughput chromosome conformation capture (Hi-C) ^3^ and chromatin interaction analysis by paired-end tag sequencing (ChIA-PET) ^4^ are some of the commonly used methods to study 3D genome organization. Hi-C detects all possible genome-wide pairwise interactions between loci. By using Hi-C maps, it was observed that chromosomes are partitioned into two compartments, A and B, representing active and inactive chromatin regions, respectively ^3^. Analysis of relatively high-resolution Hi-C maps (~40 Kbp) resulted in the discovery of self-interacting genomic regions called topologically associating domains (TADs) ^5–8^. Much higher resolution Hi-C maps (in the range of 1 – 5 Kbp) have revealed enhancer-promoter contacts ^9^.

Hi-C identifies all chromatin contacts but does not specify the proteins associated with 3D interactions. This is partially addressed by including a chromatin immunoprecipitation (ChIP) step with the Hi-C protocol. For example, ChIA-PET captures genome-wide interactions associated with specific proteins. ChIA-PET has facilitated the discovery of chromatin interactions associated with transcription factors (ER, AR), RNA Polymerase II, and structural proteins such as the cohesin component RAD21 and CTCF ^10–12^. However, Hi-C and ChIA-PET experiments are labour-intensive, time-consuming, and expensive ^9,13^. Furthermore, there always exists a possibility that the experiment outcome may not be of the desired quality. The ENCODE portal has provided RAD21 ChIA-PET datasets for about 24 cell lines ^11^. However, we still do not have the RAD21 ChIA-PET for many other cell lines ^14,15^. We still do not fully understand the key determinants of cohesin-mediated chromatin interactions. Therefore, we sought to develop a machine learning method to predict cohesin-associated chromatin interactions using simple 2D chromatin and other associated genomic features.

Machine learning has been applied to solve long-standing questions in biology. Notably, the AlphaFold system has been applied to accurately predict the 3D shape of a protein from its amino acid sequence ^16^. Several machine learning systems have been developed to understand 3D genome organization ^17–31^. For instance, transcription factor and histone modification ChIP-Seq data were used to predict the chromatin interactions between loop-associated ERa binding sites (laERBSs) ^17^. Higher-order chromatin organization A/B compartments, originally calculated using Hi-C data ^3^, have been predicted from epigenetic data, such as DNA methylation microarray, DNase hypersensitivity sequencing, single-cell ATAC sequencing, and single-cell whole-genome bisulfite sequencing ^18^. In ^22^, the authors developed a neural network to predict chromatin structural types (i.e., to which subcompartments ^9^ the chromatin loci belong) from ChIP-Seq signals. They used the available ENCODE ChIP-seq data for the GM12878 cell line (84 protein binding and 11 histone modification experiments). They have also trained a reduced model using only the 11 histone modification experiments ^22^. Moreover, Gradient Boosting regressor was used to predict the interaction frequency between loci of 25 Kbp size (the model was shown to work also at 5 Kbp resolution) ^25^. In the final model, RNA-seq data, CTCF binding, and orientation were used as the regression model predictors ^25^. In Chromatin Interaction Neural Network (ChINN), DNA sequences of interacting loci were used to predict CTCF-, RNA polymerase II- and Hi-C-associated chromatin interactions ^28^. However, most of the existing models for predicting chromatin interactions are binary classifiers and do not predict interaction strength. In addition, most of them restrict the predictions to enhancer-promoter interactions and restrict the distance between interacting loci to a few megabase pairs. Computational methods to predict the strength of cohesin-mediated cell-type-dependent interactions in a genome-wide manner are still unavailable.

In this study, we present Chromatin Interaction Predictor (ChIPr), a suite of regression models based on DNN (DNN-ChIPr), random forest (RF-ChIPr), and gradient boosting (GB-ChIPr), respectively, to predict the strength of chromatin interactions between any two anchor peaks. Our main assumption is that the interaction strength between any pair of peaks depends on a set of factors that can be easily measured or widely (publicly) available. We hypothesized that the interaction strength between two peaks depends on (A) the enrichment of the protein of interest in the two peaks (feature 1), which can be measured by ChIP-Seq, (B) the enrichment of active and inactive histone modifications (features 2 and 3), which can also be measured by ChIP-Seq, and (C) additional factors that can be easily calculated without any new experimental data, like the genomic distance between the two peaks, the GC content of the two peaks, and the CTCF motif orientation in the two peaks (features 4 to 6). These six features were selected as inputs for our model. The output of ChIPr is the predicted strength of the interaction between any two peaks/regions of interest.

We demonstrate that the predictions of ChIPr correlate well with the original ChIA-PET (as our positive control) interactions at the peak-level resolution and bin sizes of 25 and 5 Kbp. We show that ChIPr accurately predicts most of the cell-type-dependent loops identified by either ChIA-PET or Hi-C. Moreover, we have analyzed the importance of each of the model inputs for the model’s prediction accuracy and performed a detailed analysis for the role of CTCF motif orientation and CTCF occupancy in the prevalence and strength of cohesin-mediated chromatin interactions. Remarkably, our results demonstrate that, with a single experimental data (RAD21 ChIP-Seq), ChIPr can predict cohesin-mediated chromatin interactions with high accuracy.

## RESULTS

### ChIPr predictions correlate well with the original data at the peak-level resolution

The schematic of the method and a few examples of the contact maps that can be constructed using the predicted outputs at different resolutions are shown in **Fig. 1A and B**, respectively. Additional details about the input features and the regression models can be found in the “Methods” section. For each of the three variants of ChIPr—DNN-ChIPr, RF-ChIPr, and GB-ChIPr—we trained two main models using the data of the two cell lines, GM12878 and K562, respectively. We chose GM12878 and K562 because they are two of the best-characterized cell lines in the ENCODE portal ^14,15^, with the highest data quality. In addition, using models trained on two different cell lines reduces the inherent biases which might be observed due to the presence of structural variations and mutations in the genome. We used the models trained on the RAD21 ChIA-PET data from GM12878 to predict RAD21 interactions’ strengths in the cell lines K562, H1, and HepG2 using the six inputs described in **Fig. 1A**—RAD21 ChIP-Seq, H3K27ac ChIP-Seq, H3K27me3 ChIP-Seq, the genomic distance between peaks, GC content, and CTCF motif orientation flag. The CTCF motif orientation flag is an input (set to 1 if CTCF motif orientations in the two interacting peaks are convergent, and is set to ‘0’ otherwise). Reciprocally, we used the models trained on the RAD21 ChIA-PET data from K562 to predict the strengths of RAD21 interactions in the cell lines GM12878, H1, and HepG2. The RAD21 ChIP-Seq data used in our studies were not derived from RAD21 ChIA-PET data and therefore represent bonafide independent datasets. We have previously shown that ChIA-PET interaction strengths follow a negative binomial distribution ^32^. Hence, to evaluate the performance of our ChIPr, we generated random values for the interactions’ strengths drawn from negative binomial distributions with the same mean and variance as that of the corresponding original ChIA-PET sample. We measured the correlation coefficient values between the predictions we obtained for the four cell lines (using the models trained on GM12878 and K562 data, respectively) and the original ChIA-PET data. We found that the predicted outputs of the three different variants of ChIPr correlated significantly better with the original data than the randomly generated interactions’ strengths (**Fig. 2A and B, Supp. Fig. 1A and B**). We also found that the three different regression models—DNN-ChIPr, RF-ChIPr, and GB-ChIPr—yielded comparable results (**Fig. 2A and B, Supp. Fig. 1A and B**). In addition, the results for the cell lines H1 and HepG2 are quite similar for the models trained on GM12878 and K562 data, respectively (**Fig. 2A and B, Supp. Fig. 1A and B**). These results showcase the accuracy, reproducibility and generalizability of ChIPr.

**Fig. 1.**
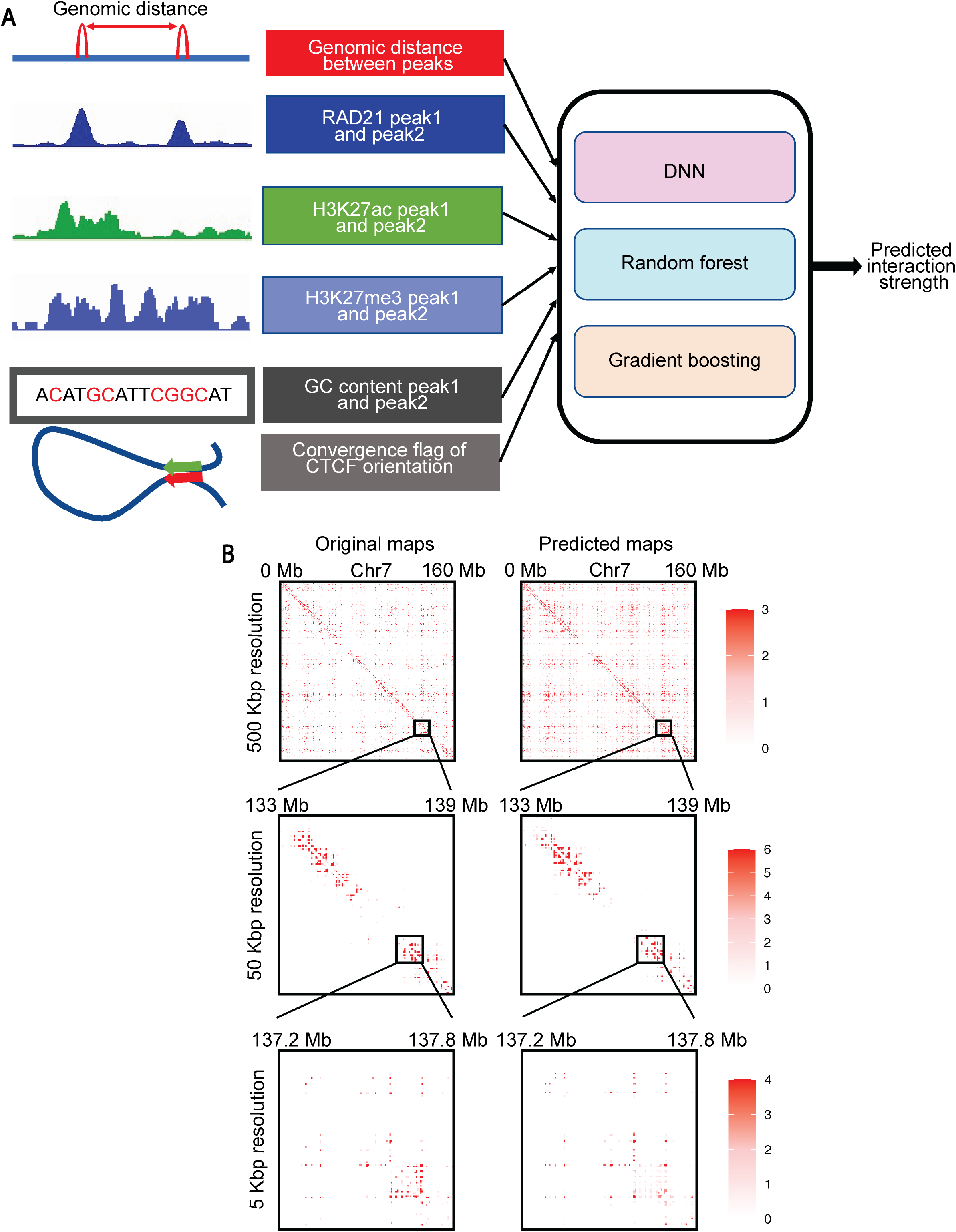
Overview of ChIPr, a regression model with three variants to predict interaction strength between two peaks. (A) A schematic representation of ChIPr showing all the input features, the three regression variants, and the expected output from ChIPr. (B) An example of contact maps constructed from the original ChIA-PET data and the corresponding ones constructed from the predictions of DNN-ChIPr at resolutions of 500 Kbp, 50 Kbp, and 5 Kbp. Heatmaps in (B) were plotted using HiTC.

**Fig. 2.**
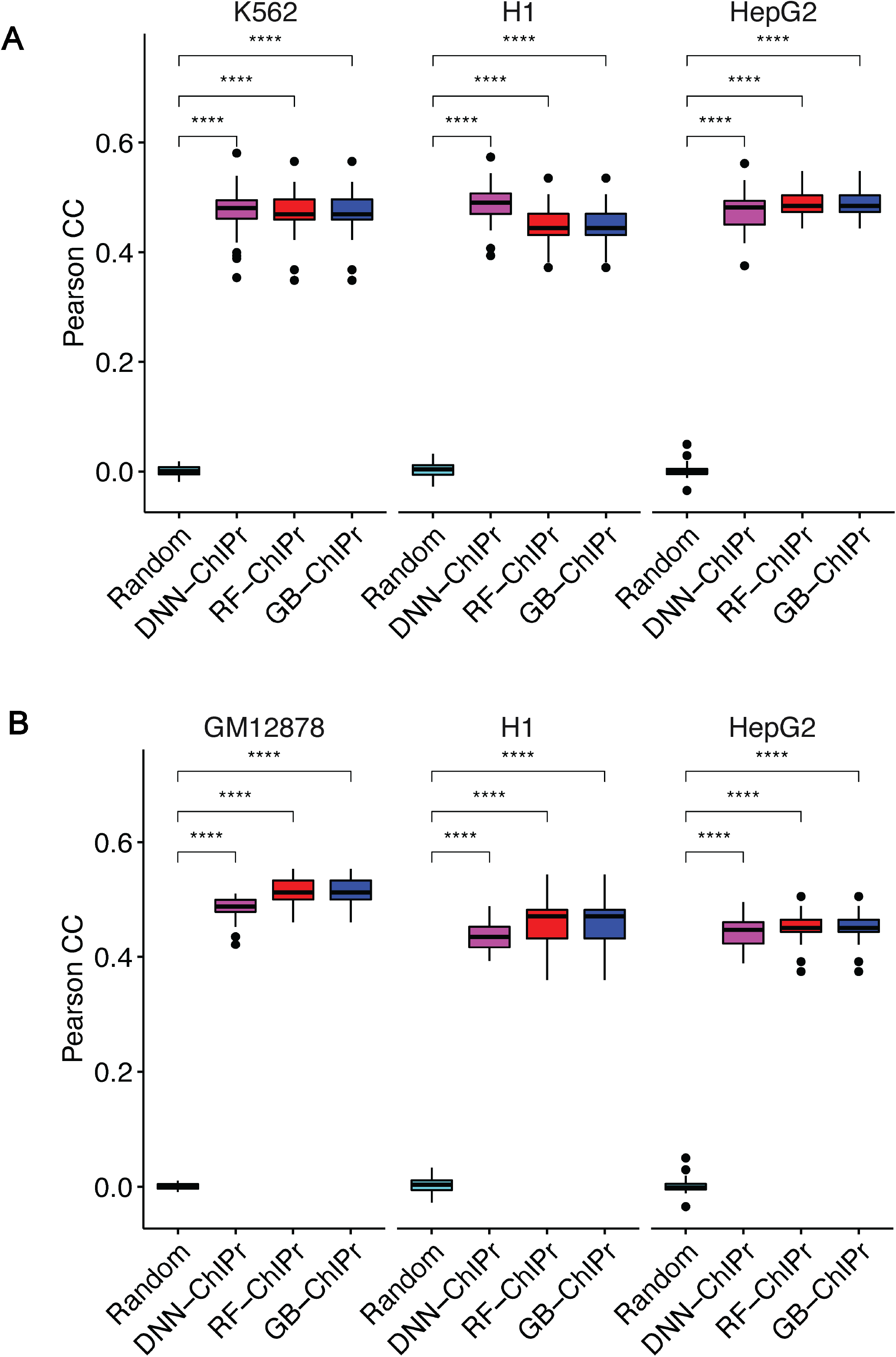
Predicted interactions correlate well with the original ones at the peak-level-resolution. (A) Predicted interactions using the three variants of ChIPr for the cell lines K562, H1, and HepG2 correlate significantly better than the random interactions with the original ChIA-PET interactions of these three cell lines. The predictions in (A) are obtained using models trained on the GM12878 cell line data. (B) Predicted interactions using the three variants of ChIPr for the cell lines GM12878, H1, and HepG2 correlate significantly better than the random interactions with the original ChIA-PET interactions of these three cell lines. The predictions in (B) are obtained using the models trained on the K562 cell line data. ****: p-value < 0.0001, Wilcoxon rank sum test.

### ChIPr predictions correlate well with the original ChIA-PET data at 25 and 5 Kbp bins resolution

Although our goal is to predict the chromatin interactions’ strengths at the peak-level resolution, we can still capture much information at lower resolutions. For instance, we can predict TADs using contact maps of 25 and 5 Kbp resolutions ^9^. Thus, we sought to measure how well the ChIPr outputs correlate with the original data at these lower resolutions.

In ^33^, HiCRep was developed to assess the reproducibility of Hi-C data taking into account its unique spatial features, such as domain structure and distance dependence. HiCRep minimizes the effect of noise by smoothing the Hi-C maps. It also addresses the impact of distance dependence by dividing the contact maps into strata. It calculates the Pearson correlation coefficient between every two corresponding strata in the two maps being compared. The weighted sum of these Pearson correlation coefficients is called the stratum-adjusted correlation coefficient (SCC). SCC has the same range and interpretation as standard correlation coefficients ^33^. In ^34^, a faster and more computationally efficient version of HiCRep was developed.

We used SCC and Pearson correlation coefficients to evaluate the similarity between the original data and the outputs of ChIPr. More specifically, we created interaction maps for the original, predicted, and randomly generated interactions at 25 and 5 Kbp bin sizes. We measured SCC and Pearson correlation between the original maps vs. the predicted and random ones. For SCC, we set the smoothing window half-size *h* to ‘2’ and the maximum genomic distance to include in calculations to 25 Mbp. We found predicted maps correlate significantly with the original maps than the random ones (**Fig. 3A-D, Supp. Fig. 2A and B**).

**Fig. 3.**
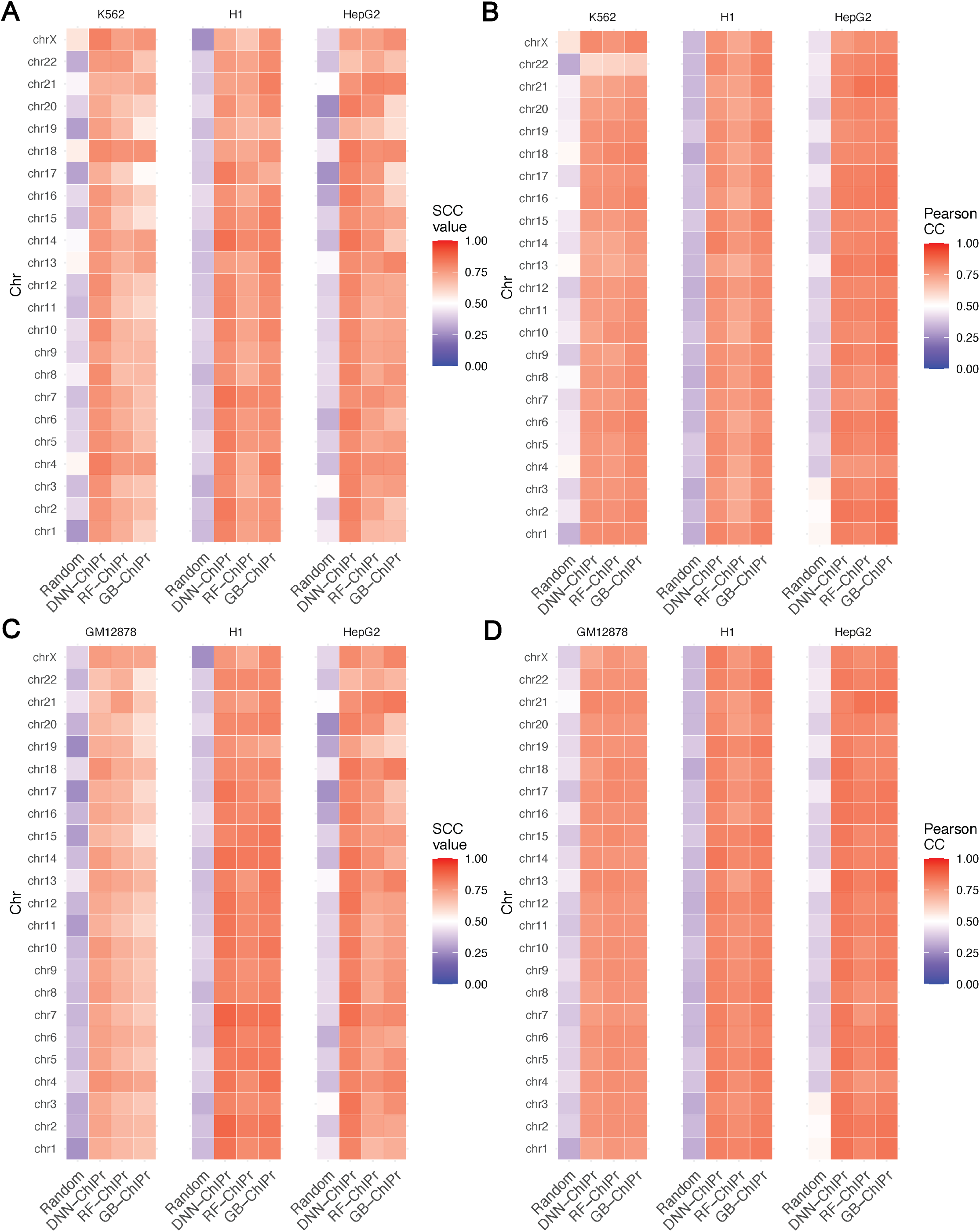
Predicted interactions correlate well with the original ones at the 25 Kbp bin resolution. (A and B) Comparison between the correlation coefficient values between the original interactions and the predicted ones using the three variants of ChIPr vs. those between the original and randomly generated ones for the three cell lines K562, H1, and HepG2. The correlation coefficients were calculated using stratum adjusted correlation coefficients (A) and Pearson correlation coefficients (B), respectively. The predictions in (A and B) were generated using the models trained on GM12878 data. (C and D) Comparison between the correlation coefficient values between the original interactions and the predicted ones using the three variants of ChIPr vs. those between the original and randomly generated ones for the three cell lines GM12878, H1, and HepG2. The correlation coefficients were calculated using stratum adjusted correlation coefficients (C) and Pearson correlation coefficients (D), respectively. The predictions in (C and D) were generated using the models trained on K562 data.

In addition, we calculated the *expected contact maps* at 5 Kbps, where each entry contains the average interaction strength at this genomic distance. We calculated the maps *P1* = (predicted/expected) for predictions obtained by the three ChIPr models and *O1* = (original/expected). We calculated the correlation between non-zero entries in *P1* and *O1.* We observed high pearson correlation values between the two matrices **(Supp. Fig. 3A-C)**. All these results show the agreement between original and predicted contact maps. This agreement highlights the ability of ChIPr to reproduce reasonably accurate contact maps with relatively small bin sizes like 25 Kbps and 5 Kbps.

### ChIPr captures ChIA-PET identified cell-type-dependent interactions

In ^11^, ChIA-PET was used to study the cohesin-mediated chromatin loops in 24 cell lines. The authors pooled ~125,000 interactions across all the cell lines and found that ~28% of that pan-cell line loop set are variable loops (i.e., cell-type-dependent loops). These variable loops are strong in certain cell types and weaker or near noise level in other types.

We investigated the whole list of cell-type-dependent loops to see if they are captured by ChIPr as strong interactions in the corresponding cell-types (i.e., interaction strength (PETs) greater than or equal to ‘3’). As a negative control, we introduced an equal number of random interactions by shuffling the coordinates of the first peak of the cell-type-dependent loops of each chromosome (see **Fig. 4A**). These randomly introduced loops were not expected to be predicted by ChIPr as strong interactions. We found that, on average, 74%, 78.5%, and 72.15% of the cell-type-dependent loops are captured in the four cell lines using DNN-ChIPr, RF-ChIPr, and GB-ChIPr, respectively (**Fig. 4B-G**). On the other hand, 2.3%, 2.6%, and 4.7% of the randomly introduced interactions were predicted as strong interactions using DNN-ChIPr, RF-ChIPr, and GB-ChIPr, respectively (**Fig. 4B-G**). These results highlight the utility of ChIPr in predicting cell-type-dependent cohesin-mediated chromatin interactions.

**Fig. 4.**
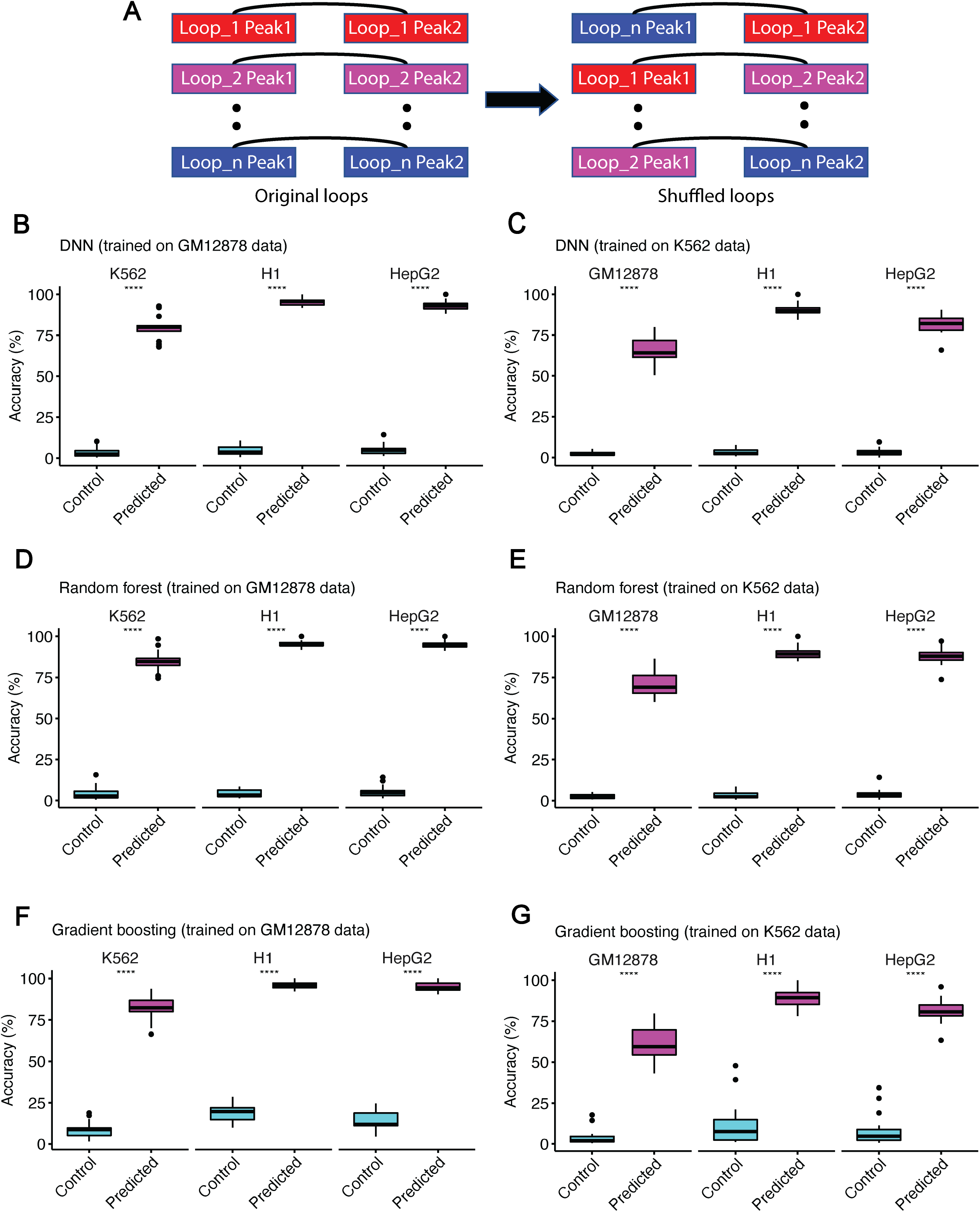
Predicted interactions capture the majority of cell-type-dependent loops. (A) An illustration of how the random control loops were generated for the comparison. (B – G) Predicted interactions using DNN-ChIA-Pr (B and C), RF-ChIPr (D and E), and GB-ChIPr (F and G) captured a significantly higher portion of the ChIA-PET identified cell-type-dependent loops vs. randomly introduced loops of the same number for the cell lines K562, H1, and HepG2. The models in (B, D, and F) were trained using the data of GM12878 cell line and the models in (C, E, and G) were trained using the data of K562 cell line. ****: p-value < 0.0001, Wilcoxon rank sum test.

### ChIPr captures both cell-type dependent and universal cohesin-mediated chromatin interactions

We further investigated a region around the *SMAD3* gene in the four cell lines GM12878, K562, H1, and HepG2. SMAD3 functions as a signal transducer in the transforming growth factor-beta (TGF-β) signalling pathway. It also transmits signals from the cell surface to the nucleus to regulate cell proliferation and gene activity ^35,36^. To visually evaluate and show the accuracy of the interaction strength predicted using ChIPr regression model, we compared the interactions from original ChIA-PET data to those predicted by RF-ChIPr model which was trained on GM12878 data (for K562, H1, and HepG2 cell lines) and K562 data (for GM12878 cell line), in *SMAD3* gene region. We found relatively dense, strongly predicted interactions for the cell lines GM12878, K562, and HepG2, which was consistent with the elevated activity of the enhancer elements in the corresponding region in these cell lines (**Fig. 5A**). On the other hand, we found few interactions in the case of H1, which was also consistent with the reduced activity of the enhancers in the region (**Fig. 5A**). Similarly, we examined another region covering the two genes *MED29* and *ZFP39. MED29* gene encodes for a protein which is a part of the mediator complex and functions in the regulation of transcription of nearly all RNA POLII dependent genes ^35,36^. On the other hand, *ZFP36* gene encodes for an RNA-binding protein involved in mRNA metabolism pathways ^35,36^. This region comprising a non-variable loop predicted strongly in all of the four cell lines was also in line with the original data (**Fig. 5B**).

**Fig. 5.**
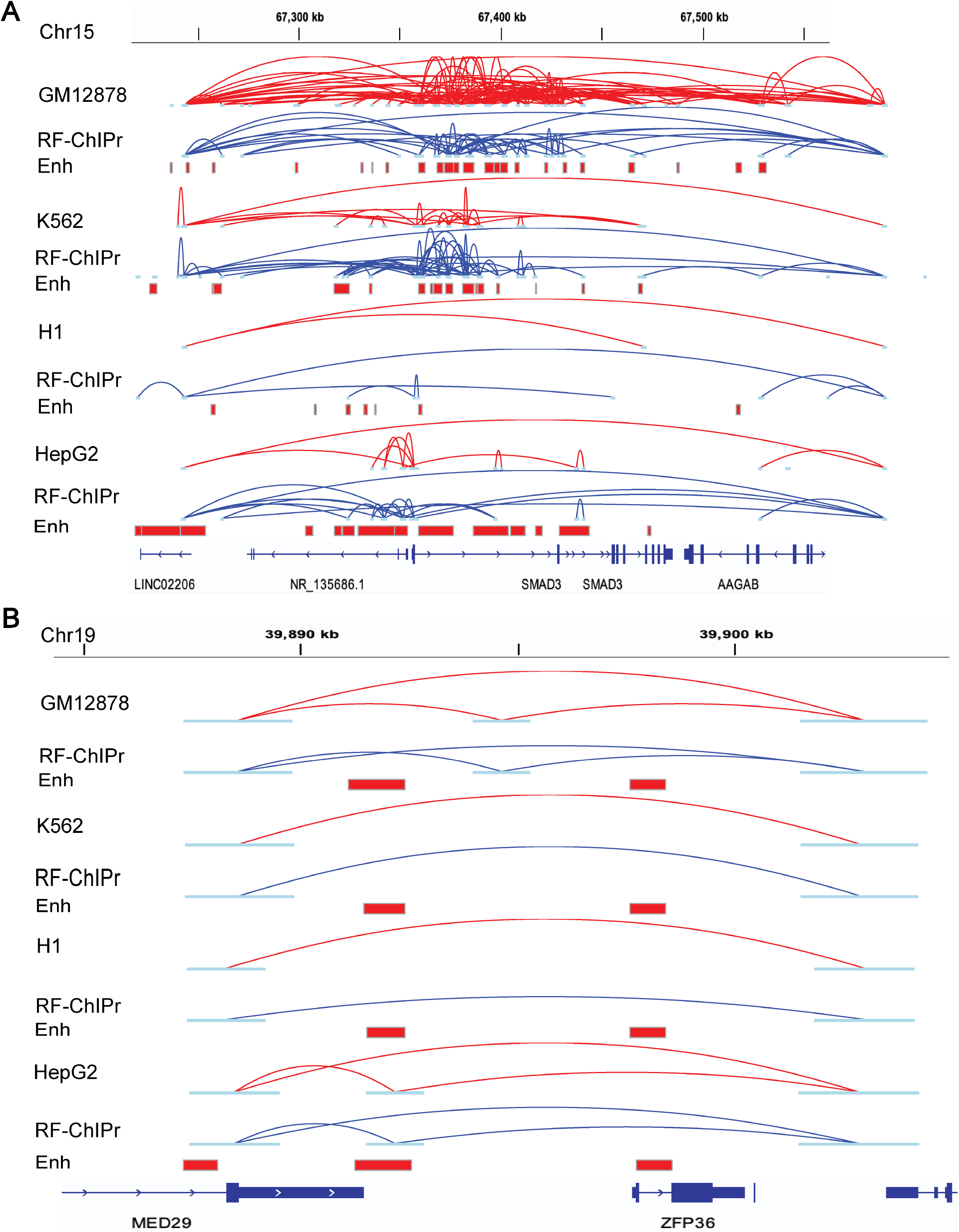
Examples for the predictions of RF-ChIPr for variable and non-variable loops. (A) Predictions of RF-ChIPr are highly similar to the original data for the selected region surrounding *SMAD3* gene. (B) Predictions of RF-ChIPr are highly similar to the original data for the non-variable loops in the region covering the two genes *MED29* and *ZFP36* in the four cell lines GM12878, K562, H1, and HepG2. Interactions shown in (A) and (B) are those having strength >=‘3’. Red: original loops from RAD21 ChIA-PET data; blue: predicted loops by RF-ChIPr.

Moreover, we also explored loops in the region surrounding the *MYC* oncogene. We found that model predictions could capture the strong interactions between *MYC* promoter and the enhancer elements located in the *PVT1* gene in the four cell lines (**Fig. 6A**). In addition, the strong set of enhancer-enhancer interactions in the regions of *CASC19* and *CASC21* genes and in the region of *PVT1* gene were also captured by all the three variants of ChIPr in GM12878 and K562 cell lines, respectively (**Fig. 6A**). We suggest that using all the three ChIPr models is likely to give a more robust view of cohesin-associated chromatin interactions in any region of interest.

**Fig. 6.**
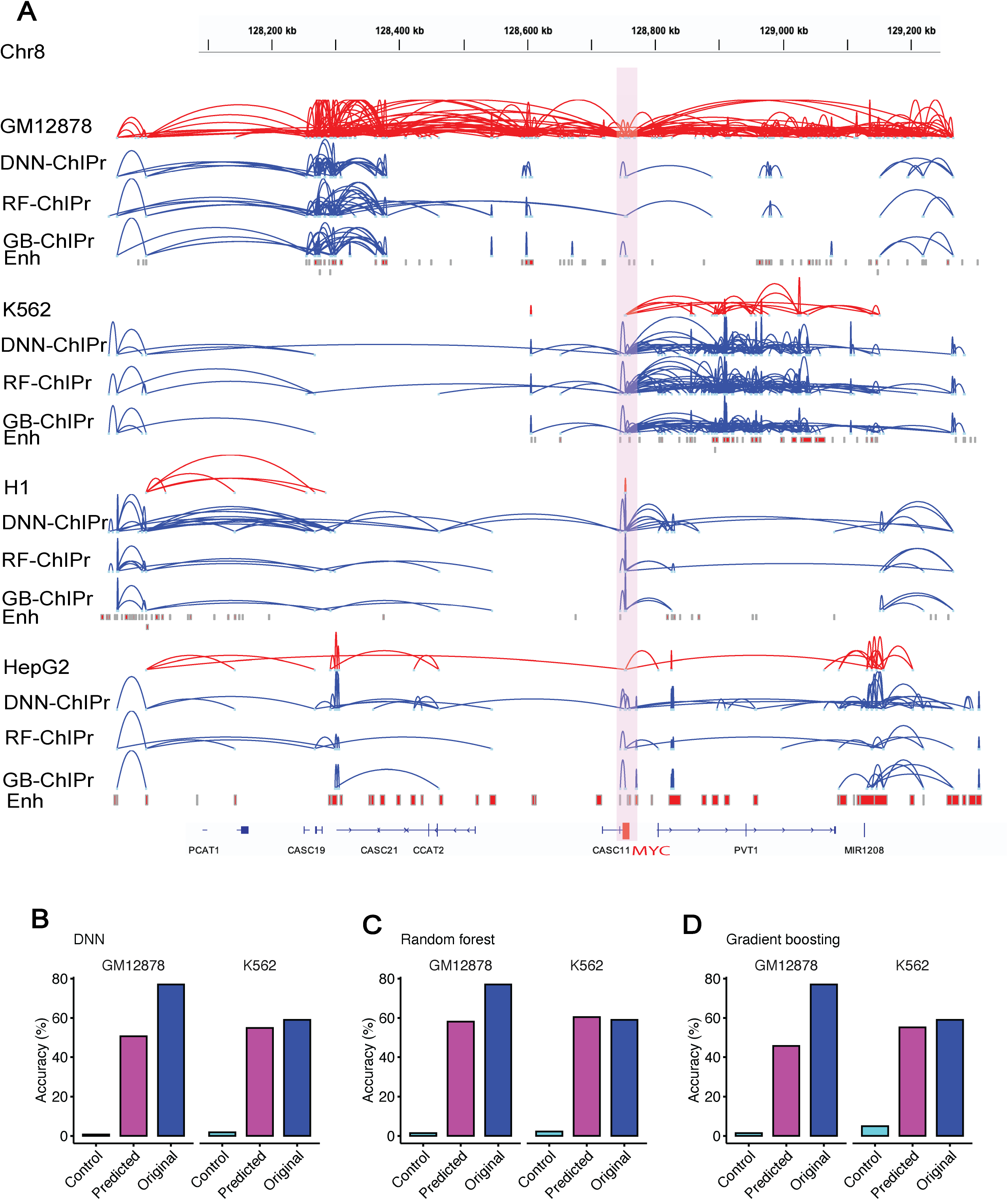
ChIPr captures majority of cell-type-dependent interactions and Hi-C identified loops. (A) Original interactions and predicted ones for the four cell lines GM12878, K562, H1, and HepG2 using the three variants of ChIPr in the region surrounding the *MYC* oncogene. Interactions shown are those having strength >=‘3’. Red: original loops from RAD21 ChIA-PET data; blue: predicted loops by ChIPr. (B - D) Predicted interactions using DNN-ChIPr (B), RF-ChIPr (C), and GB-ChIPr (D) captured the majority of the Hi-C identified loops captured by the original ChIA-PET data from the cell lines GM12878 and K562.

### ChIPr captures Hi-C identified cell-type-dependent interactions

In ^9^, in situ Hi-C was used to investigate the 3D structure of genomes of nine cell types. In addition, HICCUPS was developed to identify loops in the Hi-C maps. As an independent validation test, we measured the overlap between the strong interactions predicted by ChIPr and the Hi-C identified loops of GM12878 and K562. As a negative control, we also introduced random loops of the same number as the Hi-C identified ones (see **Fig. 4A**). We found that the predictions of our regression models capture the majority of the loops captured by the original ChIA-PET data **Fig. 6B-D**). We have also found that the Hi-C identified loops captured by the predictions of the three variants of ChIPr are significantly higher than the percentage of randomly introduced loops captured (**Supp. Fig. 4A-C**). These results suggest that a substantial number of Hi-C loops in these cell types are mediated by cohesins.

### Contributions of input features to the ChIPr predictions

To measure the importance of each input feature to the prediction accuracy, we trained the DNN-ChIPr model multiple times using the GM12878 data of odd chromosomes, eliminating one of the input features each time. We tested the trained model each time on the data of the even chromosomes and measured the performance according to the mean absolute error value when compared with the original interactions at the peak-level resolution. Then, we calculated the drop in performance when removing each of the input features (**Fig. 7A**). We found the largest drop in the performance was due to genomic distance. Hence, we concluded that genomic distance is the most important of the six input features (this is consistent to the previous ER loop predictor ^17^. We also observed an inverse relationship between RAD21 chromatin interaction strength and genomic distance (**Fig. 7B**). The second most important feature is the interaction mediating protein RAD21 ChIP-Seq data. Training the model without the H3K27ac, H3K27me3, the GC content of the two interacting peaks, or the CTCF motif orientation flag yielded a very small difference. However, when we removed both H3K27ac and H3K27me3 ChIP-Seq data together, this yielded a slightly bigger drop in performance (**Supp. Fig. 5**). This shows that, although H3K27ac and H3K27me3 ChIP-Seq signals are anti-correlated, one should use at least one of them in the training of the model. For RF-ChIPr and GB-ChIPr, we used the permutations test (see Methods section for more details), and it yielded comparable order of feature importance as for DNN-ChIPr (**Fig. 7C and D**). These results suggest that training a minimal model with a single experimental data (RAD21 ChIP-Seq data) can produce good-quality prediction results.

**Fig. 7.**
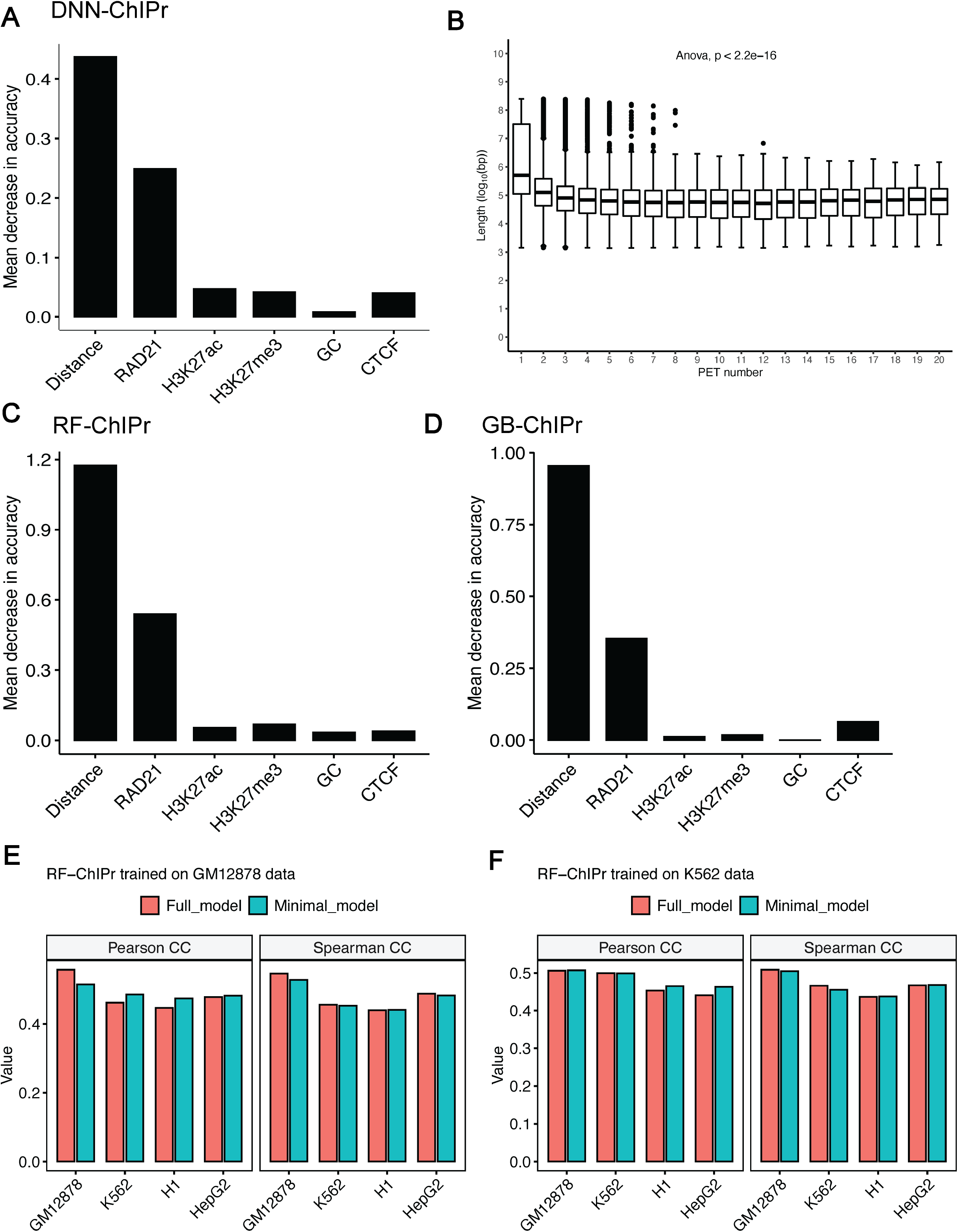
Contributions of input features to ChIPr outputs. (A) The drop in mean absolute error when comparing predicted interactions with the original ones when training DNN-ChIPr while removing one of the input features at each time. (B) The relation between the number of RAD21 interactions with different strengths and the genomic distance between the two interacting peaks. (C and D) The importance of the inputs features for RF-ChIPr (C) and GB-ChIPr (D) using the permutations test. (E and F) Comparison between the genome-level performance of RF-ChIPr minimal and full models trained on GM12878 data (E) and K562 data (F), respectively. The data is split into training data (75%) and test data (25%). In (E), the performance of GM12878 is measured on the GM12878 test data. Similarly, in (F), the performance of K562 is also measured on the K562 test data.

### Minimal model with a single experimental data—RAD21 ChiP-Seq

We tested the utility of training a minimal model using only a single experimental data—RAD21 ChIP-Seq. We trained the three regression models (DNN, Random forest and gradient boosting) with just four input data—RAD21 ChIP-Seq, genomic distance between peaks, GC content and CTCF motif orientation flag. We compared the genome level performance of the minimal ChIPr model vs. standard six input model (full model). Both models gave comparable results **(Fig. 7E and F, Supp. Fig. 6A-D)**. We also compared the performance of the minimal model with the full model by analyzing the *MYC* locus. Remarkably, both the models performed equally well in predicting the cell-type dependent cohesin-mediated chromatin interactions in the *MYC* locus **(Fig. 8)**.

**Fig. 8.**
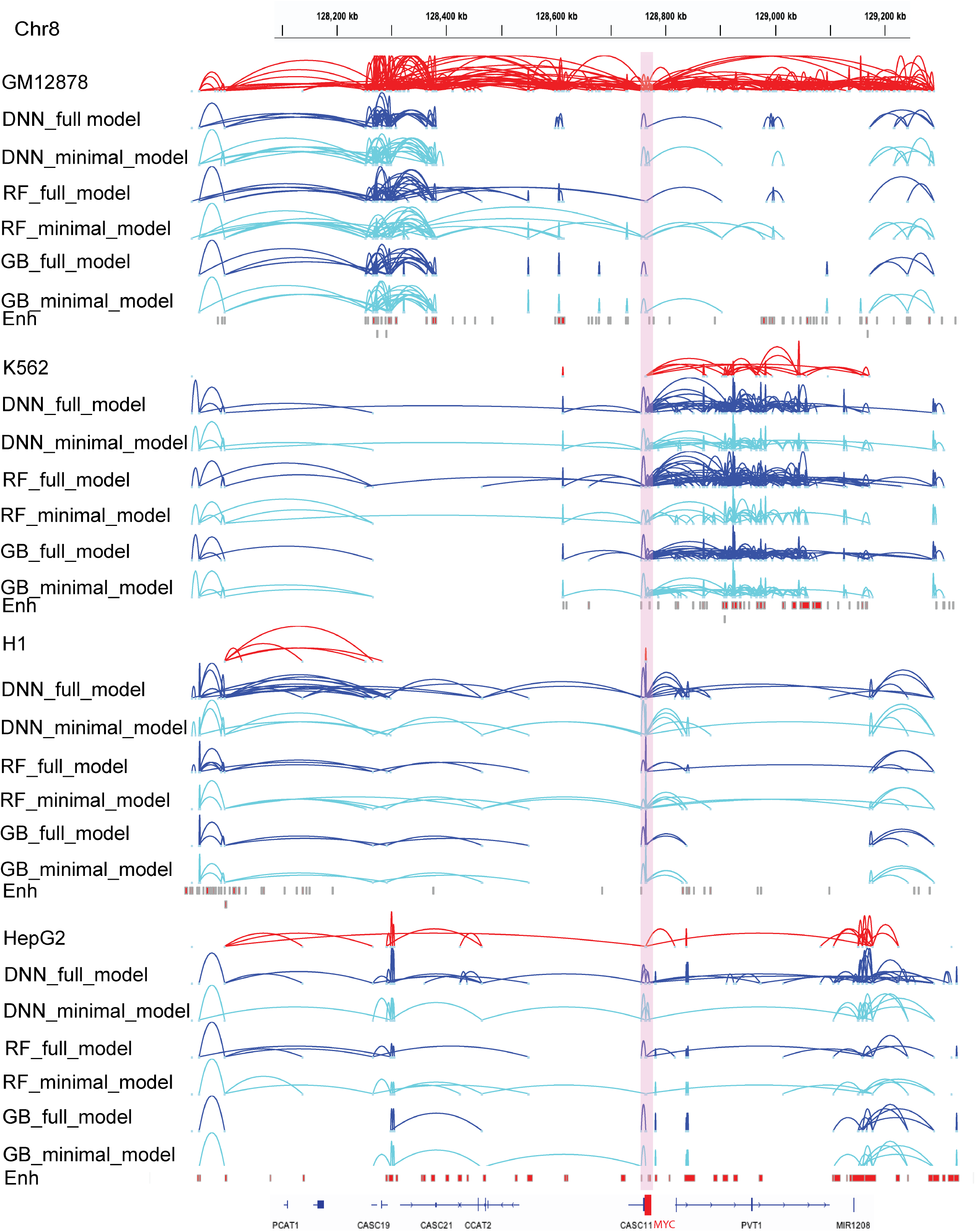
Original interactions and predicted ones for the four cell lines GM12878, K562, H1, and HepG2 using the three variants of ChIPr in the region surrounding the *MYC* oncogene. Interactions shown are those having strength >=‘3’. Red: original loops from RAD 21 ChIA-PET data; blue: predicted loops by ChIPr full model, and cyan: predicted loops by ChIPr minimal model.

### The role of CTCF motif, its orientation and CTCF occupancy in cohesin-mediated chromatin interactions

We analyzed the relationship between the strength and prevalence of the RAD21 interactions with the CTCF motif presence and orientation in the two interacting peak regions in both GM12878 and K562 cell lines. We found that the CTCF motif is found with high confidence (q < 0.3) ^37^ in both of the two interacting peaks in ~10% of the RAD21-mediated interactions in the two cell lines **(Fig. 9A)**. In addition, when the CTCF motif is present in both of the two peaks and its orientation is in the convergent manner, the interactions are, on average stronger than in the other cases, including divergent, tandem left or right, and absence of the motif in one or both peaks (**Fig. 9B and C**). A big portion of the loops with convergent CTCF motifs (45% and 34% in GM12878 and K562, respectively) exhibit strong interactions (**Fig. 9D**). However, more than 50% of the interactions are weak (PETs < 3) even with CTCF motif convergent orientation (**Fig. 9D**). On the other hand, when the CTCF motif orientation is not convergent (divergent, tandem left or right, or the motif does not exist in one or both of the two peaks), we found that more than 70% of the interactions are weak (**Fig. 9D**). These results show that the convergent CTCF motif orientation is not critical for the strength of the majority of RAD21-mediated interactions, in line with its small contribution to predicting the output of ChIPr (**Fig. 7A, C, and D**).

**Fig. 9.**
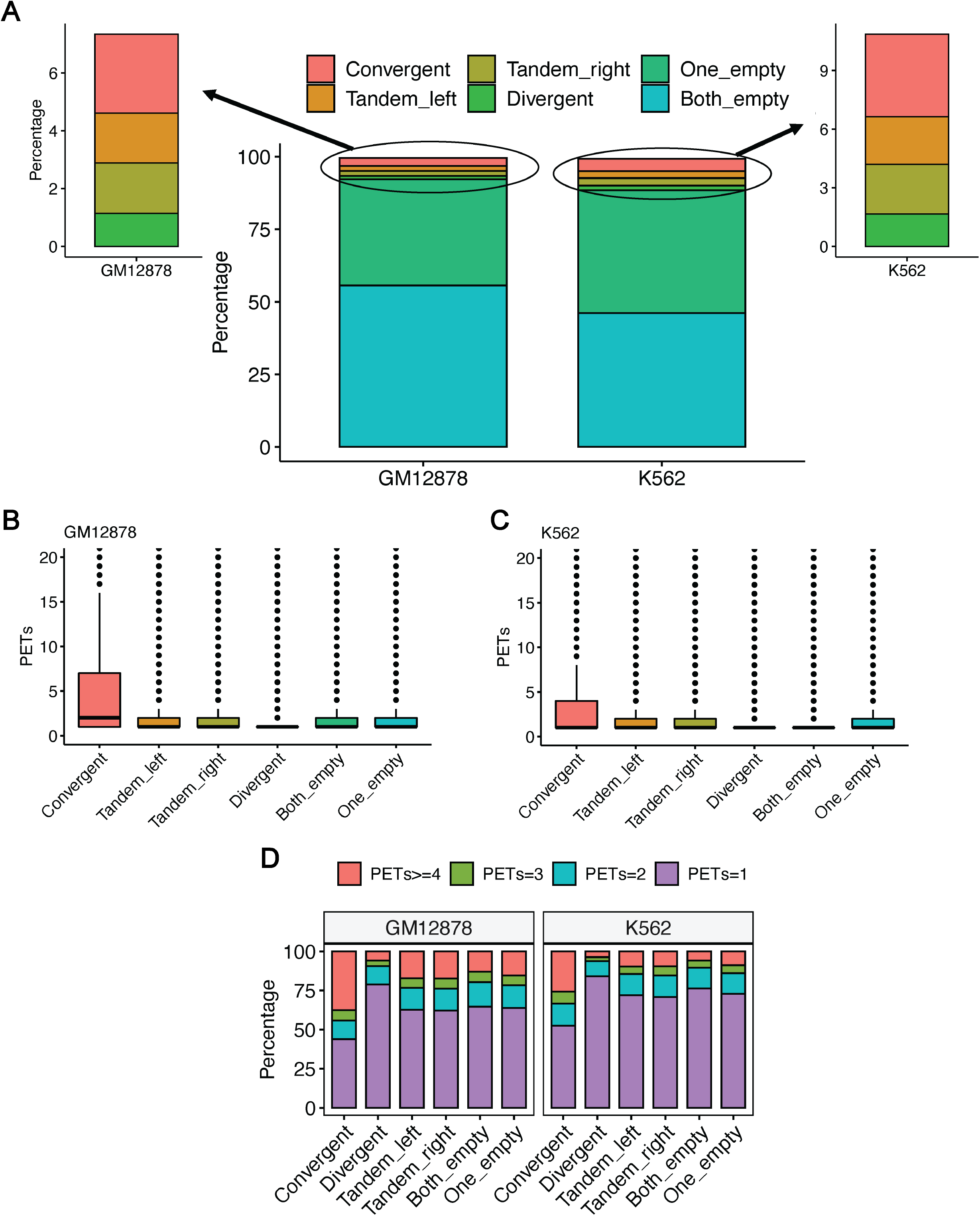
Relationship between RAD21 interactions and CTCF motif orientation and ChIP-seq binding. (A) The relationship between the RAD21 interactions prevalence and the presence of CTCF motif in the interacting peaks. (B and C) The relationship between the strength of RAD21 interactions and the presence of CTCF motif in the interacting peaks for (B) GM12878 and (C) K562, respectively. (D) The percentage of interactions of different strengths and their relation with the presence of CTCF motif in the interacting peaks.

In addition, we analyzed the relationship between RAD21 interactions and CTCF ChIP-seq peaks. This analysis showed that ~50%-80% of the RAD21 interactions were enriched with CTCF binding in the two anchor peak regions of the interaction. However, less than 15% of the interactions had no CTCF ChIP-seq binding in both of the two peaks. The peaks of the interactions with no CTCF ChIP-seq binding were enriched with enhancers, and many of these interactions were enhancer-enhancer interactions (**Supp. Fig. 7**). Taken together, these results suggest that CTCF motif presence is not a common feature of all cohesin-mediated chromatin interactions. However, CTCF occupancy is a common—but not a universal feature—of cohesin-mediated chromatin interactions. There can be multiple explanations for the discrepancy between the CTCF motif and CTCF occupancy in cohesin-mediated chromatin interactions. There could be weak or variant CTCF binding sites below our motif detection level. Indeed when we performed motif enrichment analysis for the peaks where CTCF binds without the presence of the CTCF motif in the GM12878 cell line using HOMER ^38^, we found that, in these locations, other variants of the CTCF motifs with several alignment mismatches are significantly enriched (**Supp. Fig. 8**). In addition, it has been shown that, in general, transcription factor binding may occur in the absence of any discernible motif instance, or it may occur at ‘hotspots’ where several factors are found together ^39^.

## DISCUSSION

In this study, we present ChIPr, three regression models based on DNN, random forest, and gradient boosting, respectively, and predict the strength of RAD21-mediated chromatin interactions at the peak-level resolution. ChIPr uses a few input ChIP-Seq samples and other easily obtainable public data for training, testing and prediction. We have shown that the most important feature for predicting a functional cohesin loop is the genomic distance (loop length), in line with previous report for predicting ER loops ^17^. The second most important feature was the ChIP-Seq data for the interaction mediating protein (which was RAD21 in all our analyses), consistent with the expected detection of ChIP-Seq peaks of the mediating protein at the interacting loci regions ^40^. However, we found much less importance for the two histone mark profiles, H3K27ac and H3K27me3. This may be due to the fact that these two histone marks are anti-correlated. Thus, the presence of only one of them is enough to get high prediction accuracy. When both of them were removed, we noticed a slightly bigger drop in the prediction accuracy in some cases. However, in general, the results were still very comparable. We also noticed a very small contribution by the GC content information of the two interacting peaks and the CTCF motif convergence flag. A detailed analysis of CTCF motif presence and orientation with the RAD21 interactions prevalence and strength indicated that CTCF motif presence is not necessary for RAD21 interactions prevalence. However, its presence and convergent orientation are associated in ~30%-40% of the cases with strong RAD21 interactions. These results suggest that CTCF motif presence and orientation play a necessary but insufficient role in RAD21 interactions’ strength. We have also observed the occupancy of CTCF in both of the two peaks in most of the RAD21 interactions. In the absence of CTCF binding, we found that many RAD21 loops are enhancer-enhancer interactions (**Supp. Fig. 6**).

We have shown that the RAD21-mediated DNA loop prediction outputs of ChIPr correlate well with the original RAD21 ChIA-PET data at the peak-level resolution. They also correlate well at the resolution of bin sizes 25 and 5 Kbp, which suggests that we can reliably use ChIPr predictions to detect TAD boundaries. We have also demonstrated that ChIPr could capture most of the ChIA-PET and Hi-C identified cell-type-dependent loops as strong interactions. Altogether, we have shown multiple lines of evidence that ChIPr could reliably reproduce much of the ChIA-PET information using a minimal number of easily obtainable features. These studies outline the general features of genome folding and open new avenues to analyze spatial genome organization in specimens with limited cell numbers.

## MATERIALS AND METHODS

### Structure of ChIPr

ChIPr is composed of three variants of regression models based on DNN, random forest, and gradient boosting, respectively. ChIPr uses six input features of the two interacting peaks to predict the RAD21-mediated interactions’ strengths. The first input feature is the linear genomic distance between the centres of the two peaks in kilobases. We have chosen the genomic distance because it is known to be a good predictor of the interaction strength, and it is usually inversely proportional with it according to both Hi-C and fluorescence in situ hybridization (FISH) experiments ^3,41^. The second input feature is the RAD21 ChIP-Seq data at the two interacting peaks. It is expected that the two peaks will be detected by the RAD21 ChIP-Seq data at the two interacting loci ^40^. In addition, we use the ChIP-Seq data at the two peaks for two canonical histone modification marks, H3K27ac and H3K27me3, which should correlate with active and inactive chromatin states, respectively ^9^.

Moreover, it is known that the human genome is organized into long (>300 Kbp), relatively homogeneous regions called isochores, which differ in their GC content ^42^. It has also been reported that 66% of the genes are present in the GC-rich and GC-richest isochores ^42^, suggesting a relation between gene distribution and the GC level. Accordingly, we sought that there may be a relation between chromatin activity (which leads to strong interactions) and GC content as well. Thus, we used the GC content of the two peaks as the following two input features to our regression model. Besides, it was reported in ^9^ that for the Hi-C identified loops whose corresponding anchor loci contain the CTCF motif, most of the motif pairs are convergent. Thus, we added an input that denotes the convergence of the CTCF motif orientation in the two peaks. This input is ‘1’ if the CTCF motif orientation is convergent. If the CTCF motif orientation in the two peaks is divergent, tandem left or right, or if the motif is absent in one or both peaks, the CTCF motif orientation input will be ‘0’.

### Hyper-parameter selection for DNN-ChIPr

To decide the architecture of DNN-ChIPr, we used grid search to determine the best number of layers, number of neurons in each layer, dropout rate, batch size, and activation function for the output layer. We have fixed another set of hyperparameters that are commonly used. For instance, we fixed the activation function for the hidden layers to be ‘relu’^43^. We have also used the ‘Adam’ optimizer ^44^ with a small learning rate of 10^-5^. We selected this small learning rate, although it will require a relatively longer training time to ensure the stability of the training process. In addition, we used a large number of epochs (750), with early stopping if no improvement in performance (using the validation mean square error metric) is observed for 50 epochs. The performance of each model was measured according to the mean squared error loss on the validation data. We found several models gave very comparable values of validation mean squared error (**Supp. Table1**). We chose our final model to have three hidden layers; each has 128 neurons, with ‘relu’ activation function for the output node (to ensure that the output is always bigger than zero) and values of 0.2 and 32 for the dropout rate and the batch size, respectively (**Supp. Table1**).

### Preparation of the training data

The ChIA-PET data of the four cell lines GM12878, K562, H1, and HepG2 was downloaded from the ENCODE project ^11^. The data was processed using the ChIA-PET2 pipeline ^45^ to get the inter- and intra-chromosomal interactions files. We focused on the intra-chromosomal interactions and for each interaction, we got the coordinates of the two anchor peaks and the interaction strength. We calculated the input features of anchor peaks of each interaction which comprise alongside the interaction strength a training example to our model.

### ChIP-Seq data normalization

We used the RAD21, H3K27ac, and H3K27me3 ChIP-Seq data of our four investigated cell lines (GM12878, K562, H1, and HepG2). We calculated the read count for each of the two anchor peaks of each loop. To account for the sample’s sequencing depth and the peaks’ sizes, we normalized the ChIP-Seq data using the reads per kilobase per million (RPKM) normalization method, described in the following few lines. We first get the ‘per million scaling factor’ by dividing the total number of reads in the chromosome by 1,000,000. Then, we divide the read count in each peak by the ‘per million scaling factor’, a step that accounts for the effect of sequencing depth. After that, we divide by the peak length in kilobases, a step that accounts for the peak length.

### Getting the GC content and CTCF motif orientation within peaks

To include sequence information into our model, we calculated the GC content for each peak, defined as the percentage of cytosine (C) and guanine (G) bases in that peak. We calculated it using the bedtools nuc function ^46^.

Also, we used GimmeMotifs and CTCF Bioconductor package ^37,47^ to detect the presence of the CTCF motif in each peak and its orientation. For the CTCF motif orientation input, we set it to ‘1’ if the CTCF motif orientation in the two peaks is convergent and ‘0’ otherwise.

### Constructing contact maps from peak-level interactions

ChIPr predicts interaction strength between peaks. The peak length is in the range of 2 Kb. It may be slightly smaller or bigger than that. From interactions between peaks, we can build bin-based contact maps. We do that by summing all interactions whose anchor peaks lie within the same bin. For instance, to build a 25 Kb bins contact map, we have a square matrix where each entry represents the interaction strength between two 25 Kb bins. To get the value of interaction strength between a pair of bins, we sum all the interactions whose anchor peaks fall within these two bins.

### Permutation test for determining feature importance

To calculate the permutation importance of a certain feature, a baseline metric (for instance, mean absolute error) is evaluated on the test data (for instance, the data of even chromosomes if the training was done using the data of odd chromosomes). Then, for the feature column that is required to measure its importance, this column is permuted, and the metric is evaluated again. The permutation importance of that feature is defined as the difference between the baseline metric and the metric obtained after the permutation of the feature column ^48^.

### Implementation details

The DNN model (DNN-ChIPr) was implemented using Keras, the Python deep learning API [https://keras.io/]. For the random forest (RF-ChIPr) and gradient boosting (GB-ChIPr) models, we used sklearn RandomForestRegressor and GradientBoostingRegressor with the default parameters, respectively ^49^.

### Datasets used

The RAD21 ChIA-PET data for the four cell lines GM12878, K562, H1, and HepG2 can be downloaded from the ENCODE portal [https://www.encodeproject.org]. The RAD21 ChIP-Seq data for the four cell lines can be downloaded from NCBI GEO: GSM935332 (GM12878 cell line), GSM935319 (K562 cell line), GSM935379 (H1 cell line), and GSM935647 (HepG2 cell line). The H3K27ac ChIP-Seq data for the four cell lines can be downloaded from NCBI GEO: GSM733771 (GM12878 cell line), GSM733656 (K562 cell line), GSM733718 (H1 cell line), and GSM733743 (HepG2 cell line). The H3K27me3 ChIP-Seq data for the four cell lines can be downloaded from NCBO GEO: GSM733758 (GM12878 cell line), GSM733658 (K562 cell line), GSM733748 (H1 cell line), and GSM733754 (HepG2 cell line). The CTCF ChIP-Seq data can be downloaded from NCBI GEO: GSM935611 (GM12878 cell line), GSM935407 (K562 cell line), GSM733672 (H1 cell line), and GSM733645 (HepG2 cell line). Enhancer lists for the four cell lines can be downloaded from EnhancerAtlas 2.0 [http://www.enhanceratlas.org/indexv2.php].

## Code availability

Source code of ChIPr with a detailed Readme file can be downloaded from https://git.biohpc.swmed.edu/s206442/chipr

## Acknowledgements

We thank Satwik Rajaram and Diego Castrillon for insightful comments and discussions. We acknowledge the funding support from National Cancer Institute (NCI)/NIH grant (R01CA245294).

## Author contributions

Conception and Design, A.A., M.Q.Z., R.S.M.; Methodology Development, A.A.; Data Acquisition, A.A., K.C., Y.G., and J.Y.; Data Analysis and Interpretation, A.A., K.C., Y.G., J.Y., M.Q.Z., and R.S.M.; Manuscript Writing, Review, and Revision, A.A., M.Q.Z., and R.S.M. with input from all authors; Study Supervision, M.Q.Z. and R.S.M.

