## Supplemental Figures and Legends for "Accurate prediction of cohesin-mediated 3D genome organization from 2D chromatin features"

#### SUPPLEMENTARY FIGURES LEGENDS

##### Supp. Fig. 1

Predicted interactions correlate well with the original ones at the peak-level resolution. (A) Predicted interactions using the three variants of ChIPr for the cell lines K562, H1, and HepG2 correlate significantly better than the random interactions with the original ChIA-PET interactions of these three cell lines according to Spearman correlation coefficient. The predictions in (A) are obtained using models trained on the GM12878 cell line data. (B) Predicted interactions using the three variants of ChIPr for the cell lines GM12878, H1, and HepG2 correlate significantly better than the random interactions with the original ChIA-PET interactions of these three cell lines according to Spearman correlation coefficient. The predictions in (B) are obtained using the models trained on the K562 cell line data. ****: p-value < 0.0001, Wilcoxon rank sum test.

##### Supp. Fig. 2

Predicted interactions correlate well with the original ones at the 5 Kbp bin resolution. (A and B) Comparison between the correlation coefficient values between the original interactions and the predicted ones using the three variants of ChIPr vs. those between the original and randomly generated ones for the four cell lines GM12878, K562, H1, and HepG2. The correlation coefficients were calculated using stratum adjusted correlation coefficients (A) and Pearson correlation coefficients (B), respectively. Predictions for cell lines GM12878, H1, and HepG2 were calculated using the models trained on K562 data. Predictions for the cell line K562 were calculated using the models trained on GM12878 data.

**Supp. Fig. 3**

High correlation between observed/expected contact maps at 5 Kbp bin resolution. (A – C) Observed/expected contact maps calculated for original data highly correlate with their corresponding maps calculated for data predicted by DNN-ChIPr (A), RF-ChIPr (B), and GB-ChIPr (C). Predictions for cell lines GM12878, H1, and HepG2 were calculated using the models trained on K562 data. Predictions for the cell line K562 were calculated using the models trained on GM12878 data.

**Supp. Fig. 4**

ChIPr predictions (DNN-ChIPr (A), RF-ChIPr (B), and (C) GB-ChIPr) capture Hi-C identified loops at significantly higher percentage than control loops. ****: p-value < 0.0001, Wilcoxon rank sum test.

**Supp. Fig. 5**

The drop in mean absolute error when comparing predicted interactions with the original ones when training DNN-ChIPr while removing one of the input features at each time. The plot shows that removing H3K27ac and H3K27me3 together causes a relatively bigger drop in performance than removing each of them alone.

**Supp. Fig. 6**

Comparison between the genome-level performance of minimal and full models of DNN-ChIPr (A and B) and GB-ChIPr (C and D). The models in (A and C) were trained on the data of GM12878 cell line. The models in (B and D) were trained on the data of K562 cell line. The data is split into training data (75%) and test data (25%). In (A and

C), the performance of GM12878 is measured on the GM12878 test data. Similarly, in (B and D), the performance of K562 is also measured on the K562 test data.

**Supp. Fig. 7**

The majority of RAD21 interactions have CTCF ChIP-seq binding in both of the two anchor peaks of the interactions in the four cell lines GM12878, K562, H1, and HepG2. The portion of interactions that misses CTCF ChIP-seq binding in the two anchor peaks is mostly enriched with enhancer-enhancer interactions.

**Supp. Fig. 8**

HOMER results for the top enriched motif (BORIS) and its top four best matches with known motifs in the locations of CTCF binding for the GM12878 cell line.

Supp. Fig. 1

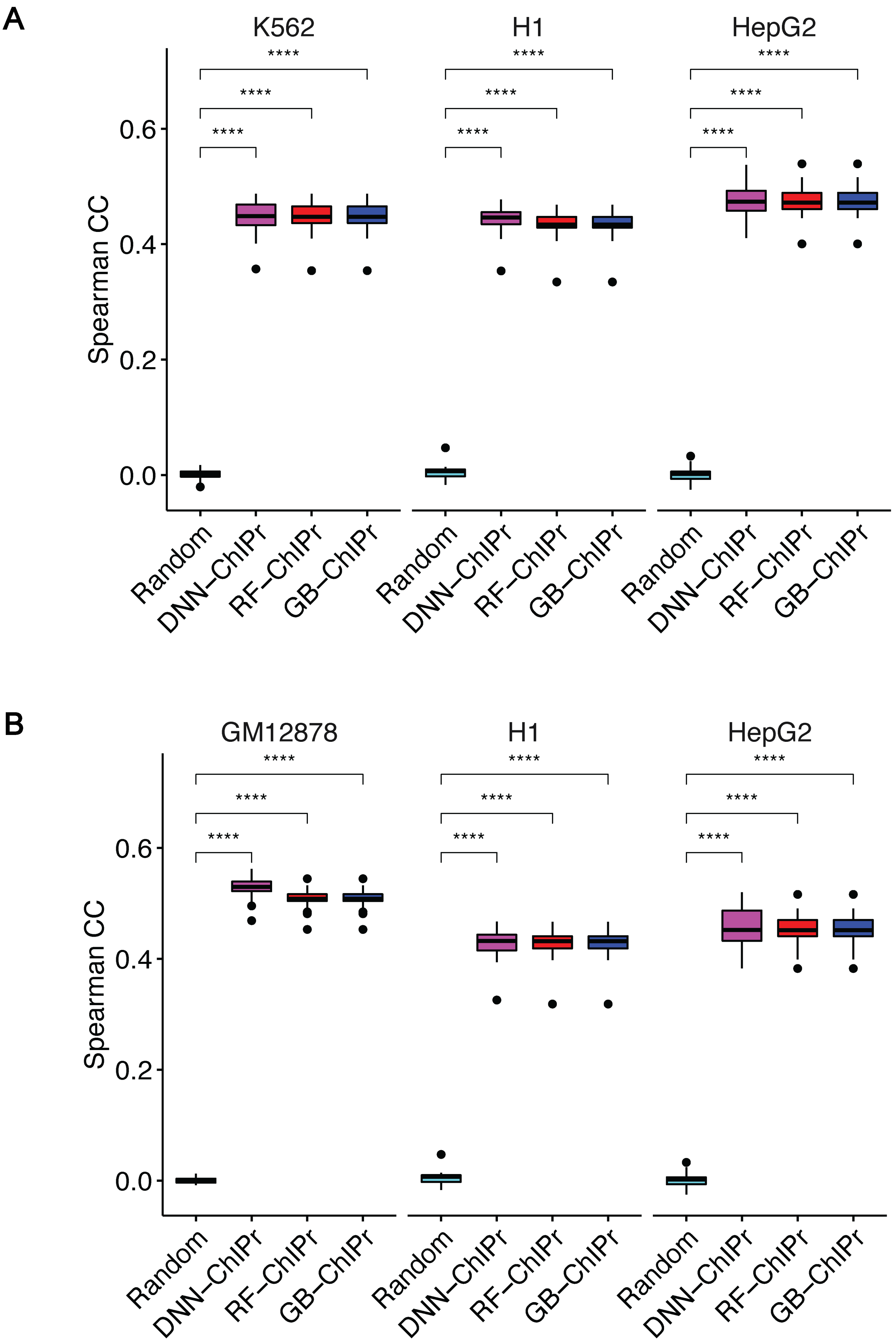
A
K562
H1
HepG2
****
Spearman CC
0.6
0.4
0.2
0.0
Random
DNN–ChIPr
RF–ChIPr
GB–ChIPr
B
GM12878
H1
HepG2

Supp. Fig. 2

A

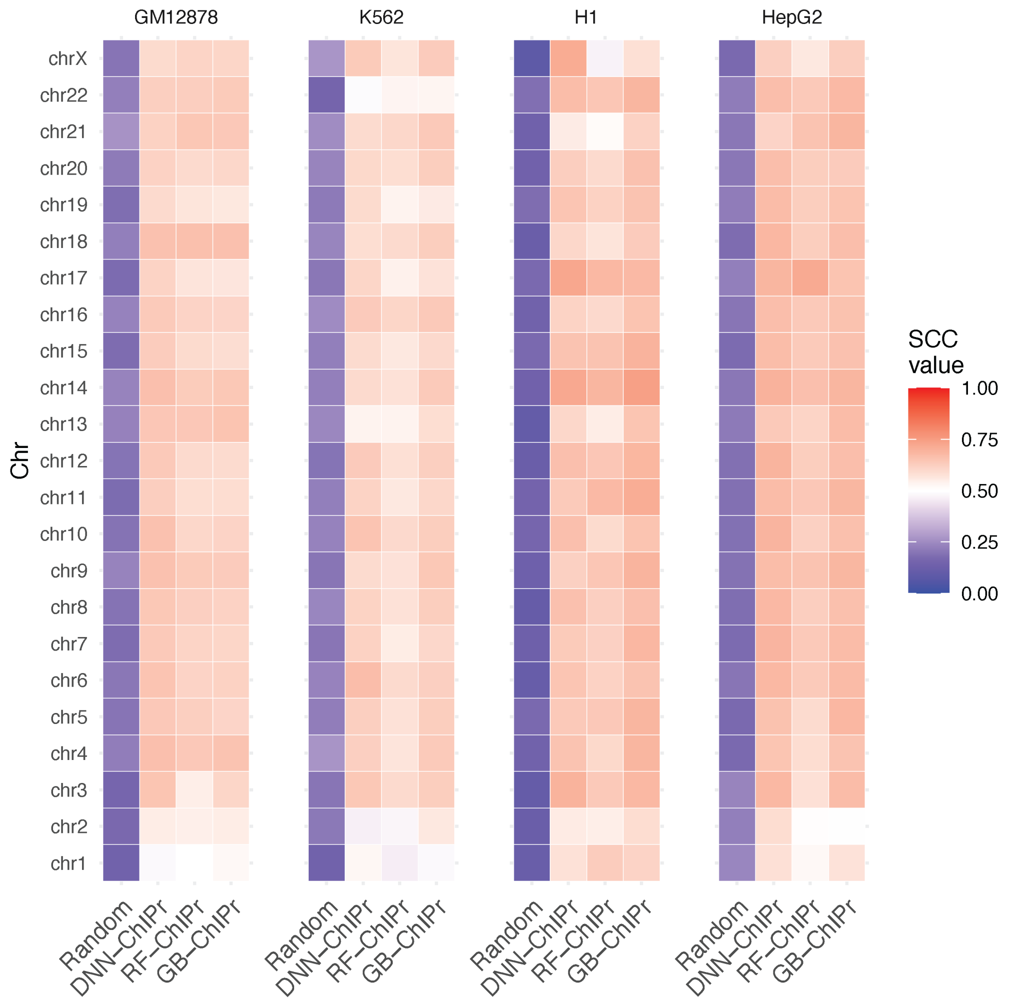

GM12878
K562
H1
HepG2
chrX
chr22
chr21
chr20
chr19
chr18
chr17
chr16
chr15
chr14
chr13
chr12
chr11
chr10
chr9
chr8
chr7
chr6
chr5
chr4
chr3
chr2
chr1
Chr
Random
DNN–ChIPr
RF–ChIPr
GB–ChIPr
SCC value
1.00
0.75
0.50
0.25
0.00


B

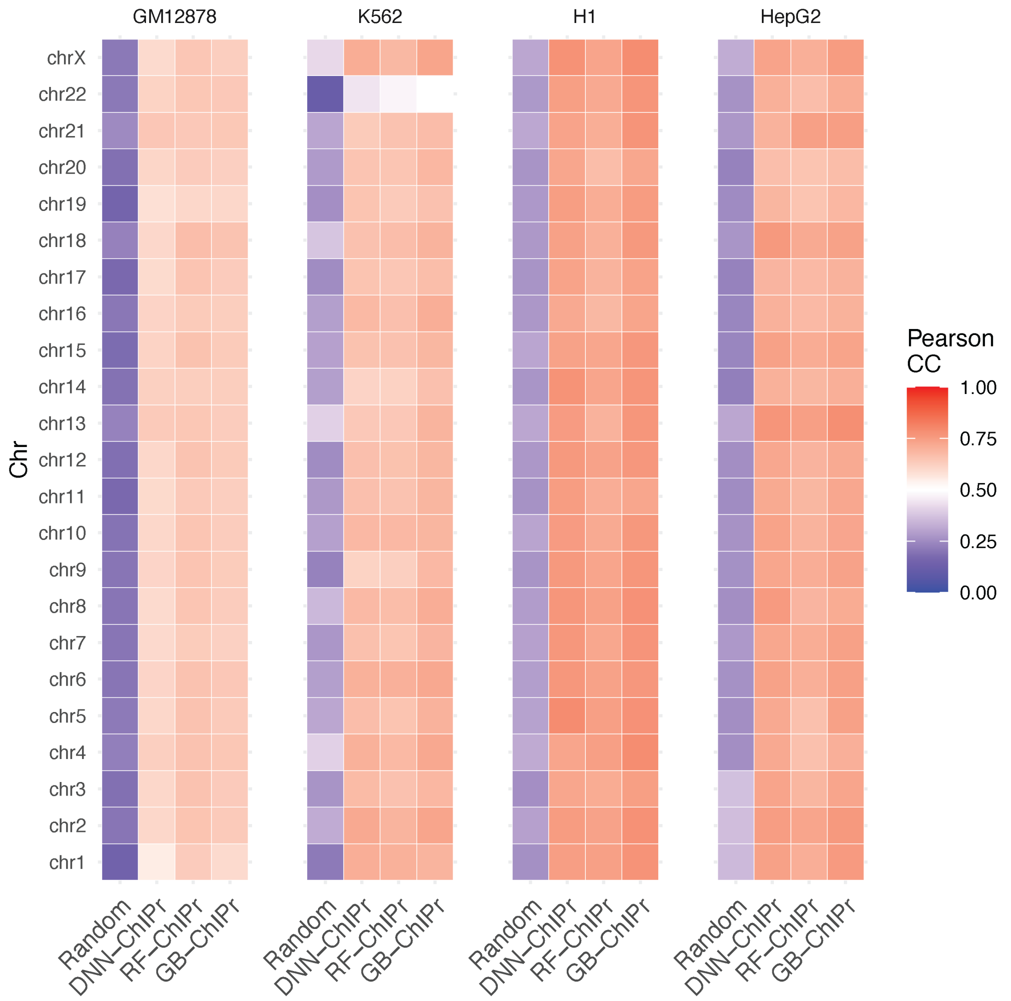

GM12878
K562
H1
HepG2
chrX
chr22
chr21
chr20
chr19
chr18
chr17
chr16
chr15
chr14
chr13
chr12
chr11
chr10
chr9
chr8
chr7
chr6
chr5
chr4
chr3
chr2
chr1
Chr
Random
DNN–ChIPr
RF–ChIPr
GB–ChIPr
Pearson CC
1.00
0.75
0.50
0.25
0.00

### Supp. Fig. 3

A

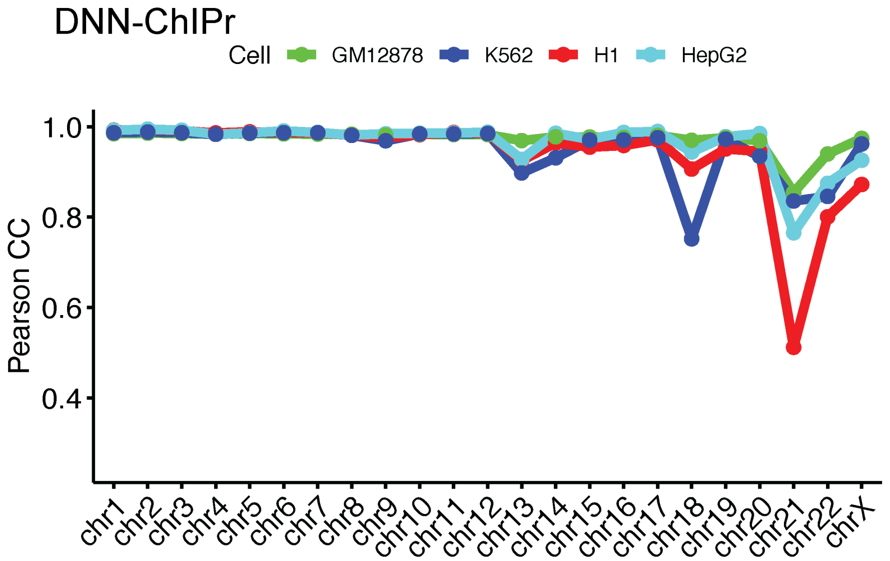


B

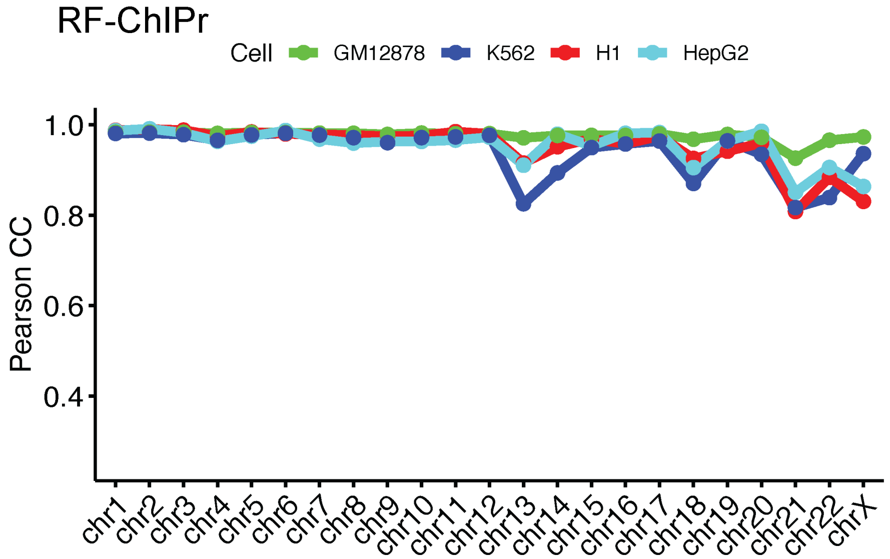


C

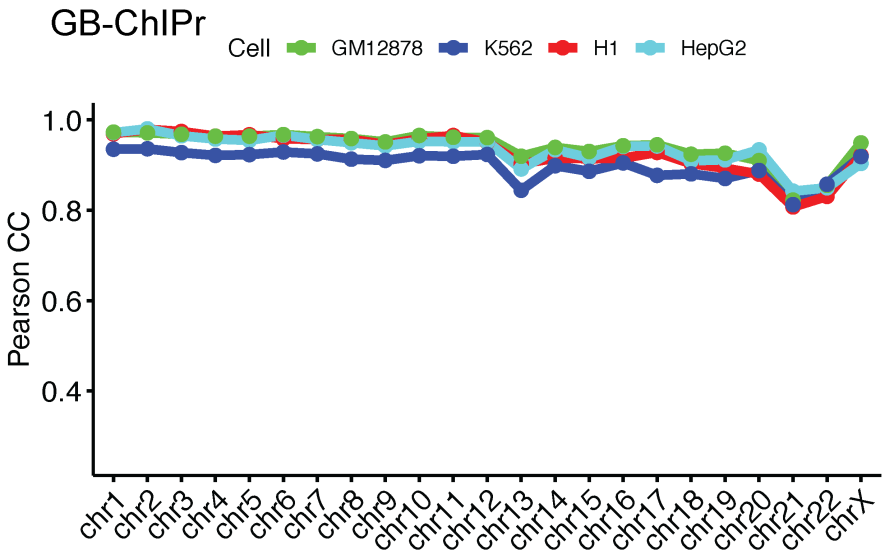

Supp. Fig. 4

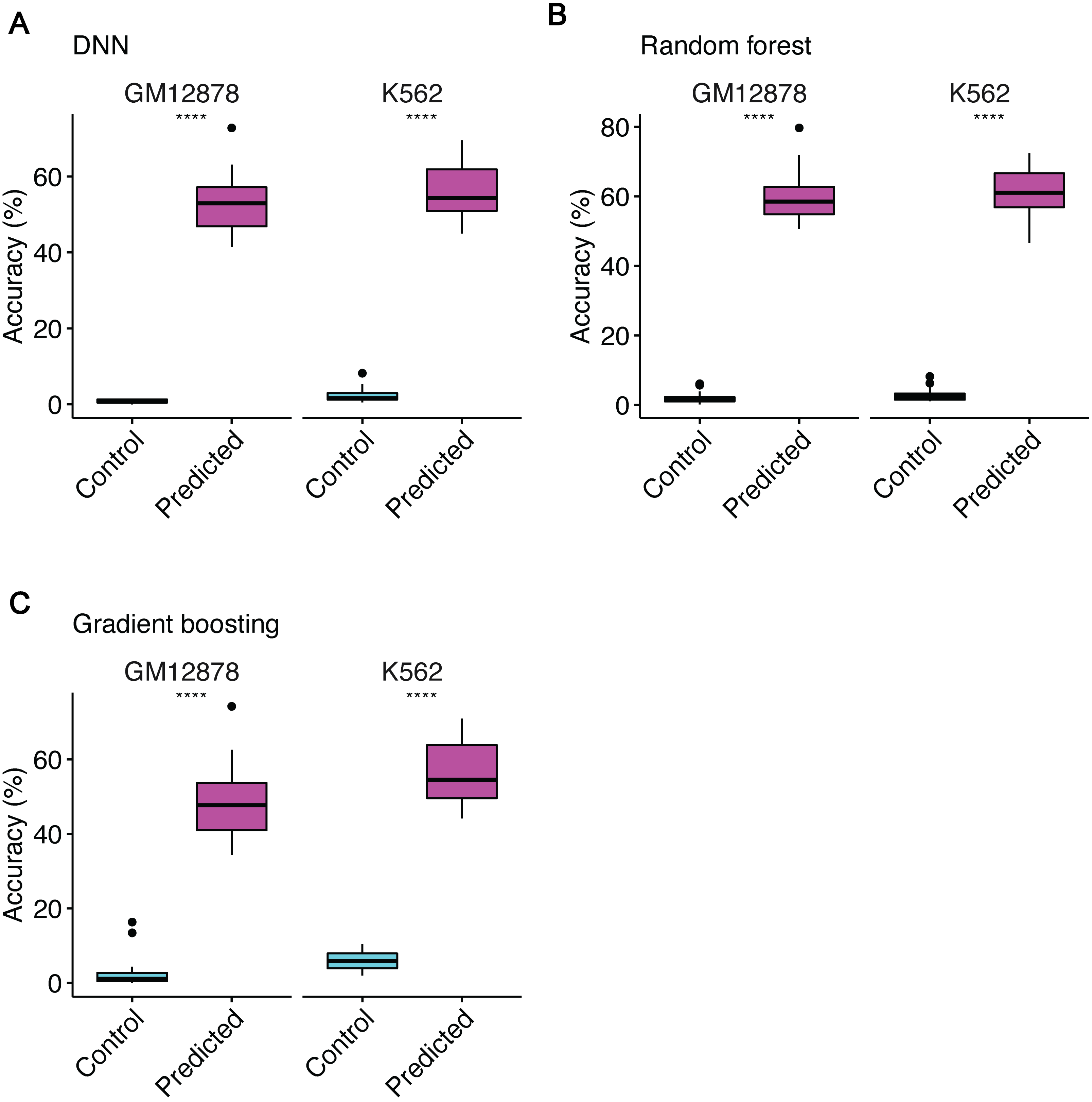

A
DNN
GM12878
K562
****
****
Accuracy (%)
0
20
40
60
Control
Predicted
Control
Predicted
B
Random forest
GM12878
K562
****
****
Accuracy (%)
0
20
40
60
80
Control
Predicted
Control
Predicted
C
Gradient boosting
GM12878
K562
****
****
Accuracy (%)
0
20
40
60
Control
Predicted
Control
Predicted

### Supp. Fig. 5

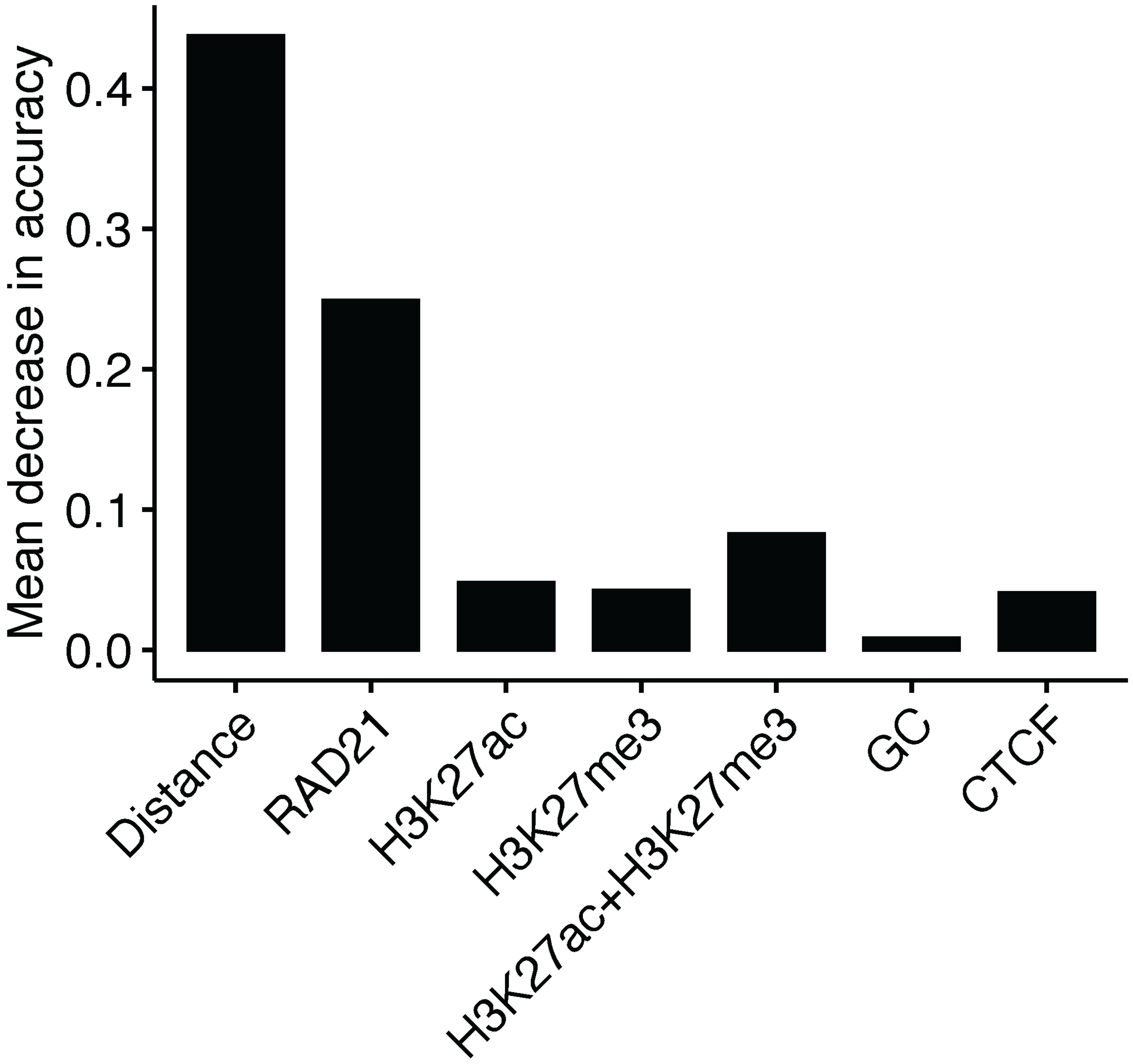

Mean decrease in accuracy
0.4
0.3
0.2
0.1
0.0
Distance
RAD21
H3K27ac
H3K27me3
H3K27ac+H3K27me3
GC
CTCF

Supp. Fig. 6

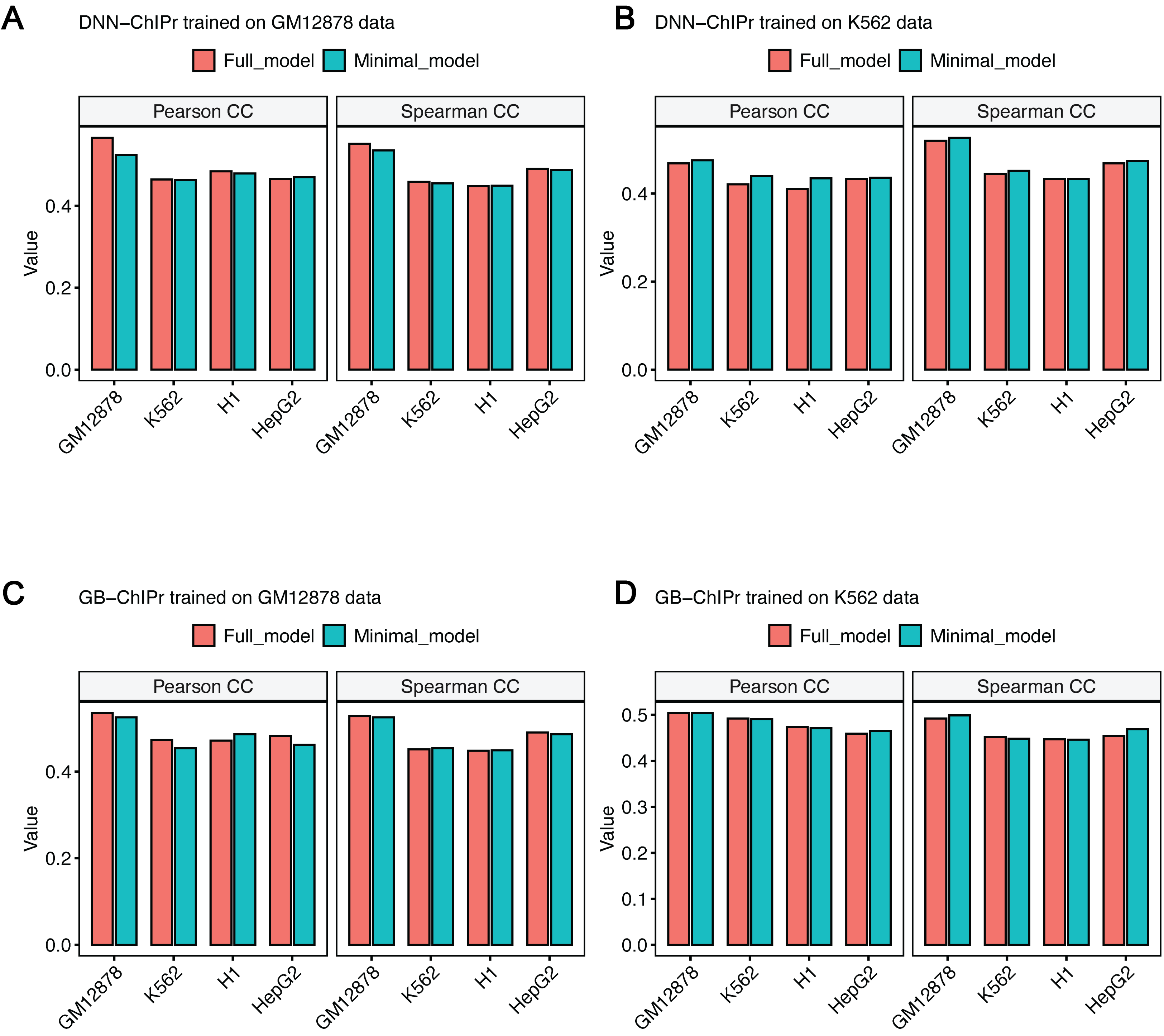
A
DNN−ChIPr trained on GM12878 data
B
DNN−ChIPr trained on K562 data
C
GB−ChIPr trained on GM12878 data
D
GB−ChIPr trained on K562 data
Full_model
Minimal_model
Pearson CC
Spearman CC
Value
GM12878
K562
H1
HepG2

Supp. Fig. 7

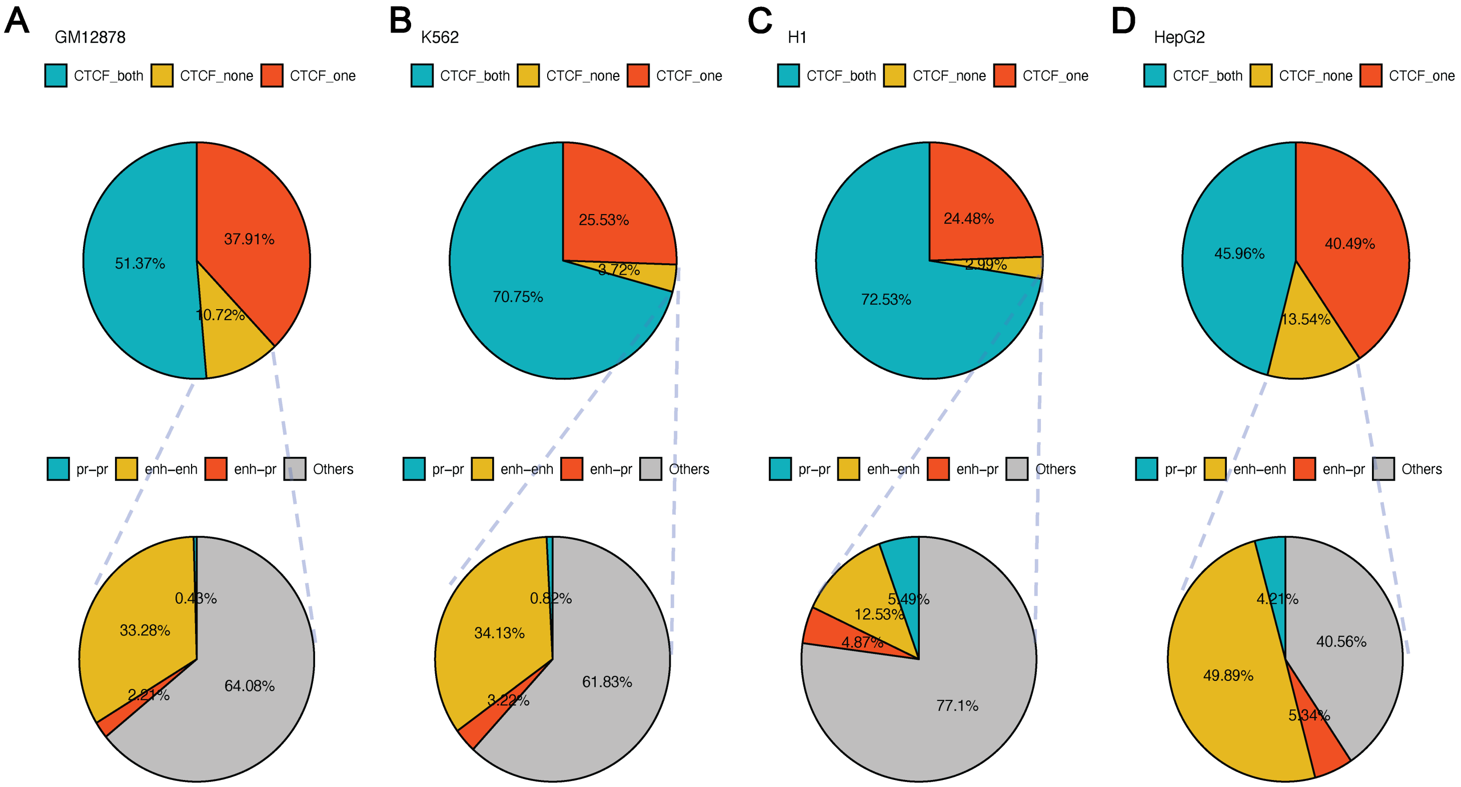

A
GM12878
CTCF_both
CTCF_none
CTCF_one
51.37%
37.91%
10.72%
pr−pr
enh−enh
enh−pr
Others
0.43%
33.28%
2.21%
64.08%
B
K562
CTCF_both
CTCF_none
CTCF_one
25.53%
3.72%
70.75%
pr−pr
enh−enh
enh−pr
Others
0.82%
34.13%
3.22%
61.83%
C
H1
CTCF_both
CTCF_none
CTCF_one
24.48%
2.99%
72.53%
pr−pr
enh−enh
enh−pr
Others
5.49%
12.53%
4.87%
77.1%
D
HepG2
CTCF_both
CTCF_none
CTCF_one
45.96%
40.49%
13.54%
pr−pr
enh−enh
enh−pr
Others
4.21%
40.56%
49.89%
5.34%

Supp. Fig. 8

**BORIS(Zf)/K562-CTCFL-ChIP-Seq(GSE32465)/Homer**

Match Rank: 1
Score: 0.91
Offset: -1
Orientation: reverse strand
Alignment: -CCACTAGRGGGC-------
GCCASCAGGGGGCGCYVNNG

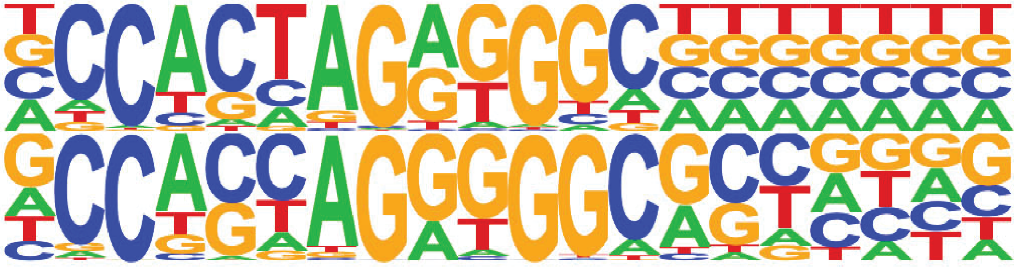

**CTCF/MA0139.1/Jaspar**

Match Rank: 2
Score: 0.90
Offset: -3
Orientation: forward strand
Alignment: ---CCACTAGRGGGC----
TGGCCACCAGGGGGCGCTA

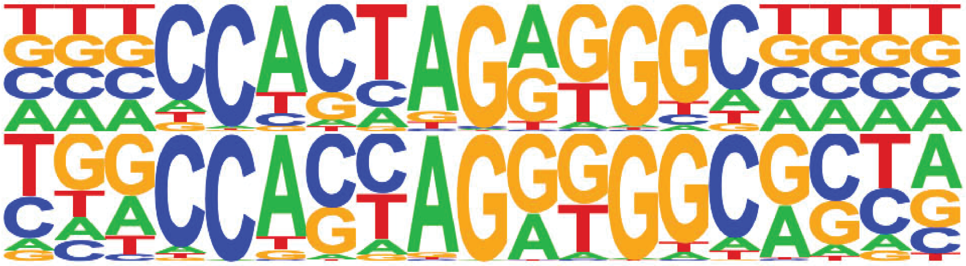

**CTCF(Zf)/CD4+-CTCF-ChIP-Seq(Barski_et_al.)/Homer**

Match Rank: 3
Score: 0.90
Offset: -3
Orientation: reverse strand
Alignment: ---CCACTAGRGGGC-----
TGGCCACCAGGTGGCACTNT

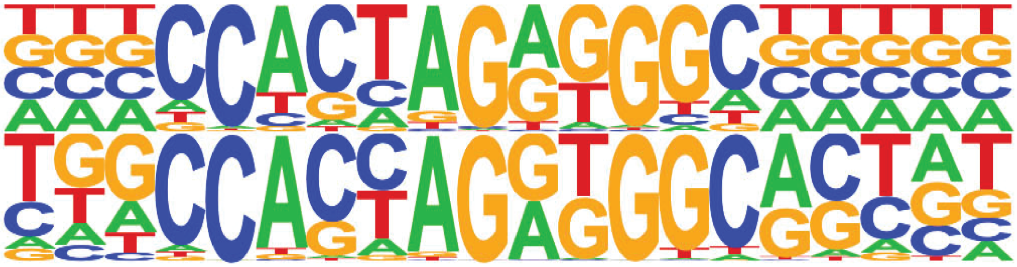

**PB0076.1_Sp4_1/Jaspar**

Match Rank: 4
Score: 0.60
Offset: 1
Orientation: reverse strand
Alignment: CCACTAGRGGGC------
-NNNAAGGGGGCGGGNNN

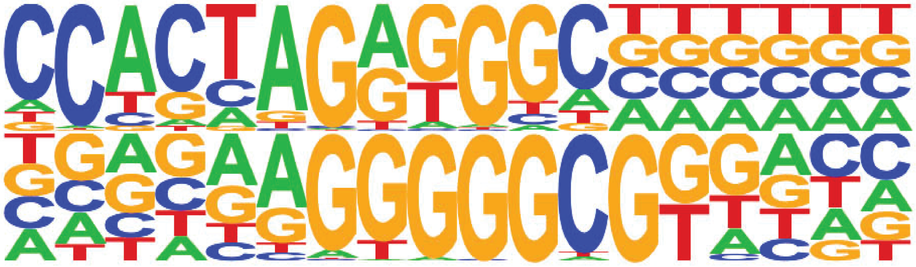
